# Pathogenic strains of a gut commensal drive systemic platelet activation and thromboinflammation in lupus nephritis

**DOI:** 10.1101/2025.06.20.641288

**Authors:** Abhimanyu Amarnani, Cristobal F. Rivera, Macintosh Cornwell, Tyler Weinstein, Susan R.S. Gottesman, Cynthia Loomis, Zakia Azad, Andy Lee, Joshua Prasad, Betsy Barnes, Mingyang Yi, Nimat Ullah, Nicolas Gisch, Kelly Ruggles, Bhama Ramkhelawon, Gregg J. Silverman

## Abstract

Imbalances in the gut microbiome have been linked to increased intestinal permeability and disease flares in systemic lupus erythematosus (SLE). Our study revealed that patients with flares of lupus nephritis (LN) and intestinal expansions of the anaerobic commensal, *Ruminococcus gnavus* (RG), displayed whole blood transcriptome profiles indicative of platelet, neutrophil, and myeloid cell activation, a profile reminiscent of sepsis. Serum analysis confirmed elevated serum levels of Platelet Factor 4 and neutrophil extracellular traps, which significantly correlated with levels of IgG-antibody to a novel lipoglycan (LG) produced by pathogenic RG strains, which was also documented in an independent LN cohort. To test for causality, *in vivo* mouse models further demonstrated that gut colonization with LG-producing RG strains, as well as a single intraperitoneal challenge with an LG preparation, caused platelet activation and megakaryocytosis in bone marrow and spleen. Mice colonized with RG strains that produce LG developed cellular infiltration of the kidneys by neutrophils and monocytes. Hence, RG expansions during renal flares may identify a specific LN flare endotype driven by thromboinflammatory mechanisms. Antibodies that arise from immune exposure to the RG lipoglycan may serve as a surrogate biomarker, helping to elucidate the impact of the relationship between gut microbiota communities and clinical outcomes in patients afflicted by LN. [208]

**Graphical Abstract:** 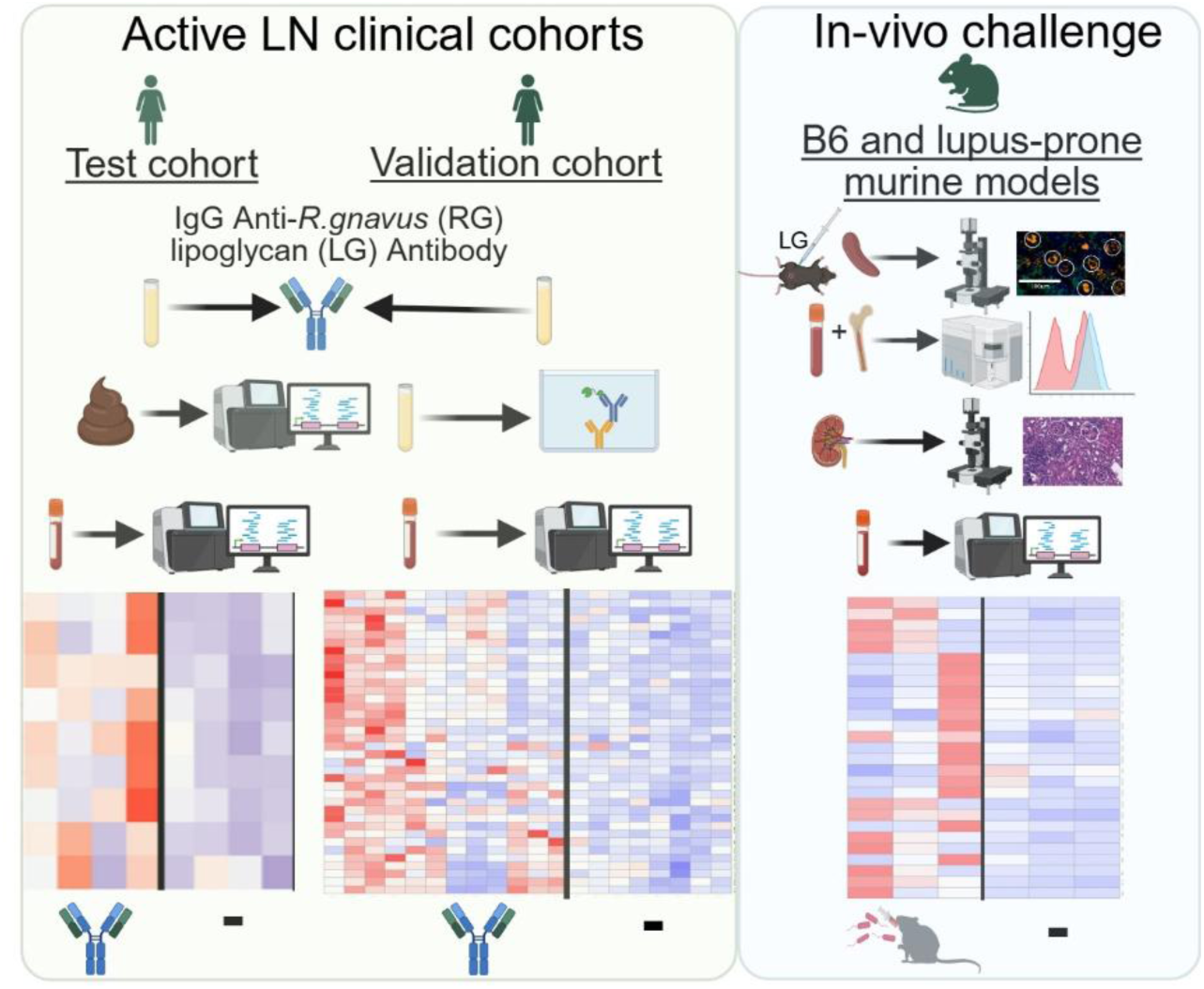

## Introduction

Systemic lupus erythematosus (SLE) is a chronic systemic autoimmune disease with great clinical heterogeneity, often marked by recurrent flares that result in severe, multi-organ injury^1^. Lupus nephritis (LN), which affects over half of all SLE patients. The pathogenesis of LN involves immune complex deposition in the glomeruli that leads to permanent glomerular and tubular damage, and despite recent advancements in therapy within ten years of diagnosis, one in five patients will progress to end-stage renal disease^2^. In genetically predisposed individuals, the influence of environmental factors on lupus pathogenesis is suspected but poorly understood^3^. We have postulated that the internal environment, posed by our gut commensals, may play key roles (reviewed in Silverman et al. 2024^4^).

The gut microbiota represents a complex community of microorganisms residing in the gastrointestinal tract, which plays crucial roles in immune development and the maintenance of immune homeostasis. Dysbiosis, or imbalances in gut microbiota, can result in increased gut permeability and systemic release of gut microbial factors, which have been implicated in diverse inflammatory and autoimmune diseases^5,6^. Renal involvement in patients with SLE, associated with gut microbial dysbiosis, has been linked to intestinal expansions of specific commensal strains of the pathobiont, *Ruminococcus gnavus* that we refer to as RG^4^ for continuity with earlier work^1,4,5,7–9^, while this species has recently been renamed *Mediterraneibacter gnavus*. A member of the *Lachnospiraceae* family of obligate anaerobes, RG is a keystone species in the gut microbiome, displaying great genomic heterogeneity between strains. In health, RG has low community representation, while it is reported to be highly expanded in patients with certain inflammatory diseases^10^. Interestingly, RG strains isolated from patients with active LN express a novel form of lipoglycan (LG). This RG-specific LG, characterized by mass spectrometry and NMR^1^, is a complex glycoconjugate with a unique structure that is attached to the bacterial cell membrane by a lipid anchor and a carbohydrate linker, with a conserved oligosaccharide core, that has extensions of one or more hexose moieties^4^. LGs produced by different RG strains have a conserved structure and are recognized by post-immunization monoclonal antibodies, as well as by spontaneously arising serum IgG anti-LG antibodies present at high levels in about one-third of LN patients^1,7^.

A pathogenic role in autoimmune disease is supported by evidence that murine colonization by lupus-specific RG strains that produce this LG toxin results in translocation out of the gut, and modulation of systemic immune responses^4^. Moreover, in lupus-prone mice, RG colonization can accelerate the progression of the autoimmune disease^11^. In lupus patients, gut expansions of RG strains with LG are associated with higher disease activity, and longitudinal studies have shown that RG blooms are temporally associated with LN flares^7^. Given these previously identified associations between intestinal expansions of pathogenic RG strains and LN flares, we sought to clarify how RG could directly influence systemic immune responses.

Our studies now show the involvement of systemic activation of platelets, which are traditionally known for roles in hemostasis and thrombosis, and in cardiovascular disease^12^. Yet, platelets are coated with receptors of the innate immune system, and they also serve as immune sentinels for the detection of invasion by microbial pathogens, triggering the secretion of inflammatory effectors. In patients with sepsis^13,14^, platelet activation leads to endothelial cell injury and promotes the triggering of neutrophils to release extracellular traps, which contributes to microthrombus formation, exacerbating septic coagulopathy and inflammatory reactions^15^. In LN, platelet activation has been implicated in end-organ leukocyte recruitment and the formation of immune complexes^12,16,17^.

Herein, studies of lupus patients in a pilot cohort, and confirmed in a validation cohort, included analyses of blood RNA sequence libraries that revealed an otherwise unexpected relationship between intestinal blooms and activation of peripheral platelets and neutrophils. Strikingly, the transcriptomic profiles in these pathways correlated with levels of serum human IgG-antibodies to the LG of RG, which arise from *in vivo* immune exposure to LG. In murine challenge models, intestinal expansions of RG strains that produce LG induced platelet activation and systemic neutrophil extracellular trap release (NETosis). RG colonization, only by strains that produce LG, induced renal deposition of NET fragments, macrophage infiltration, and renal injury. Hence, from two independent LN patient cohorts and several murine models, we provide evidence that pathogenic strains of RG promote platelet activation and thromboinflammation in patients with LN, thereby defining a distinct endotype of LN disease.

## Results

### LN patients can be dichotomized based on RG intestinal blooms

In recent studies, we reported an association between high disease activity and expansions of the anaerobic gut commensal, RG, in lupus patient cohorts^1,7,8^. In a longitudinal study, based on ACR/EULAR 2019 criteria^18^, we identified a cohort of patients with the diagnosis of SLE, that included eight patients without clinical evidence of renal disease, and eight who fulfilled clinical criteria for LN (Figure 1A) with renal biopsy documentation of WHO histopathologic class III and/or IV proliferative glomerulonephritis (GN), as well as elevated serum anti-dsDNA autoantibodies and hypocomplementemia^7^.

**Figure 1.**
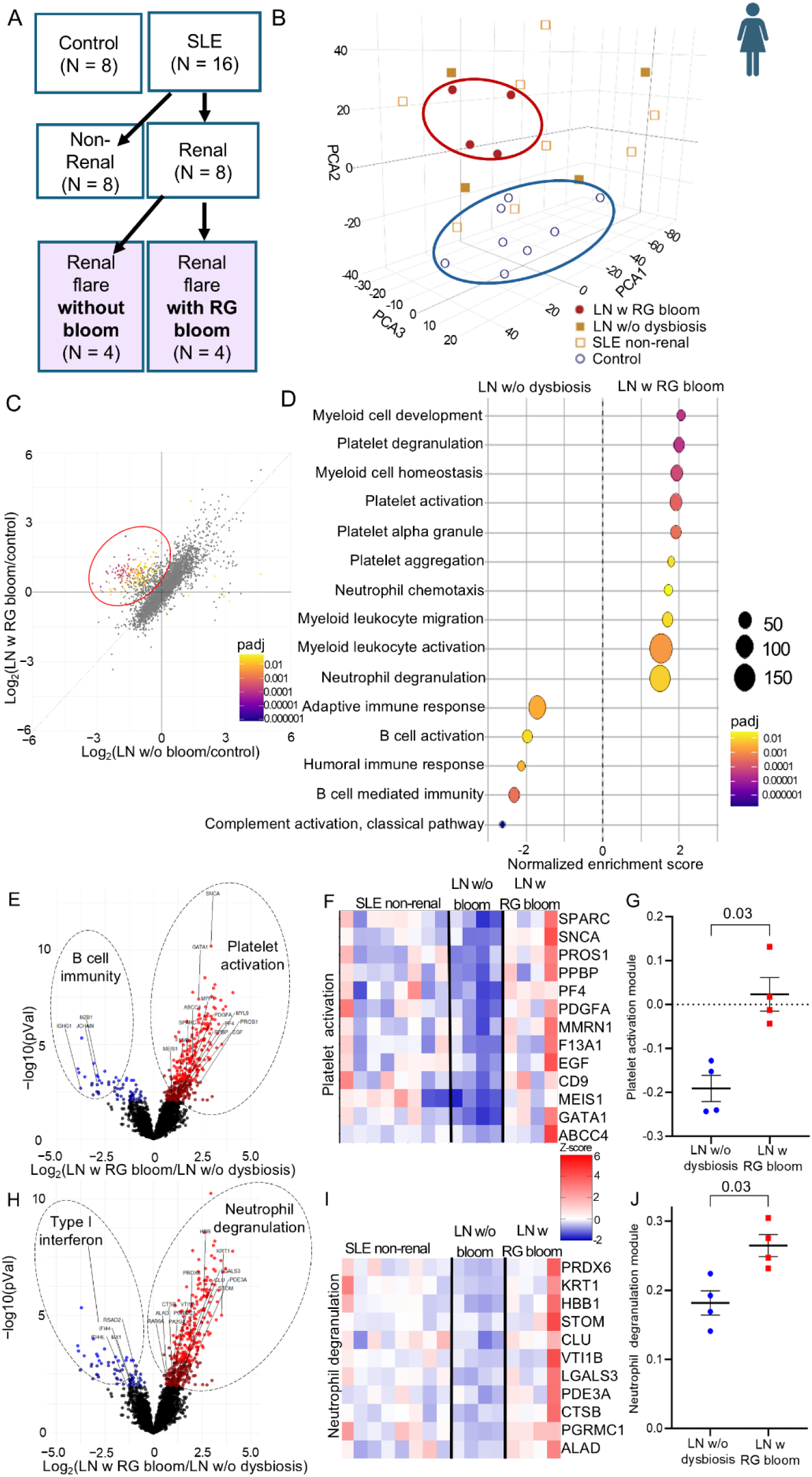
Whole-blood RNA sequence analysis of patients with LN demonstrates increased platelet activation in patients with *R. gnavus* intestinal blooms. (A) Patients with LN (all class III and/or IV) in the pilot cohort were dichotomized based on levels of anti-LG serum antibodies. (B) Principal component analysis (PCA) of gene expression in whole-blood RNAseq libraries with top 25% variance among control, SLE without LN (SLE non-renal), LN without *R. gnavus* bloom (LN without dysbiosis), and LN with *R. gnavus* gut bloom (LN with RG bloom). PCA1 40.9%, PCA2 19.1%, and PCA3 7.8%. (C) Scatter plot of Log2-fold change (gene expression) comparing LN patients with RG blooms (LN with RG bloom) vs. controls to LN patients without RG blooms vs. controls. 217 genes are significantly differentially expressed between LNRG and LN without bloom (Waldtest, qvalue<0.05), with 202 upregulated and 15 downregulated. (D) GSEA pathway analysis results of ranked genes based on LNRG vs. LN without bloom, showing significantly overrepresented pathways from genes increased in LNRG (right) or instead increased in LN without bloom (left). Myeloid cell development (NES=2.05, padj=3.2E-5), platelet degranulation (NES=1.9; padj=3.2E-5), myeloid cell homeostasis (NES=1.94, padj=7.9E-5), platelet activation (NES=1.93; padj=2.3E-4), platelet alpha granule (NES=1.93; padj=3.8E-4), neutrophil chemotaxis (NES=1.70; padj=2.6E-2), myeloid leukocyte migration (NES=1.70, padj=1.2E-2), myeloid leukocyte activation (NES=1.55, padj=1.5E-3), and neutrophil degranulation (NES=1.5; padj=7.8E-3). B cell activation (NES=-1.8; padj=2.9E-2), humoral immune response (NES=-2.6; padj=1.4E-7), and complement activation (NES=-2.1; padj=5.7E-3). (E) LN with RG bloom is devoid of B cell activation and classical complement pathway; instead, these pathways are associated with LN without bloom. Gene expression of LNRG vs. LN without bloom, with genes related to platelet activation (right) and B cell activation (left), noted. (F) Heatmap of gene expression in platelet activation in SLE non-renal LN, LN without bloom, and LN with RG patients. (G) Singscore module assessment of platelet activation pathway in LN patients with and without gut dysbiosis. (H) Gene expression of genes related to neutrophil degranulation (right) and Type I interferon (left), with (I) heatmap of gene expression in SLE non-renal, LN without bloom, and LN RG patients, and (J) Singscore module of neutrophil degranulation genes in LN patients with and without gut dysbiosis. Unpaired, non-parametric t-test.

Of the LN patients identified with previously reported criteria^1,7^, four of these patients were found to have RG intestinal expansions associated with high-level serum antibody to the RG lipoglycan (LG), while the remaining LN patients had neither such anti-microbial antibody elevations nor detected expansions of individual gut taxa compared to healthy subjects^1,7^. The clinical features of these LN patients were heterogeneous for the presence of rash and arthritis, and there were no significant observed differences based on anemia, lymphopenia, neutropenia, platelet counts, or serum albumin^7^. Patients were treated with hydroxychloroquine alone, or in combination with azathioprine or mycophenolate mofetil, and received doses of less than 40 mg of prednisone daily (**Extended data table 1**).

### Differential blood transcript expression in LN patients with RG blooms

To provide an unbiased perspective on the state of the immune system in each SLE subject, we isolated RNA from whole blood in RNA-stabilizing tubes in one batch (see methods). For the analysis of the RNAseq libraries, unsupervised principal component analysis (PCA) was then performed. Based on differential expression between each subject group, we identified the top 25% expressed genes. As expected, there was a clear transcriptomic distinction between SLE patients and the control subjects (**Figure 1A, Extended data**). Surprisingly, compared to patients *with nonrenal disease and the* LN patients without gut dysbiosis, *we* discovered differences in the libraries of LN patients with RG blooms (**Figure 1A, B, Extended data 1**).

We next assessed gene expression in each of the groups, and compared the gene expression profiles of LN patients with RG blooms versus controls, and in LN patients without RG blooms versus controls (**Figure 1C**). This analysis demonstrated a set of 217 significantly differentially expressed genes, with 202 upregulated in LN patients with RG blooms compared to LN patients without blooms. The other 15 genes were relatively under-expressed. To further define the significance of these differentially expressed genes based on function, we completed gene set enrichment analysis (GSEA) of genes with a greater than two fold change (log2FC) between LN patients with RG blooms versus LN patients without RG blooms, and an adjusted p-value <0.05. Of all pathways significantly enriched in this analysis, 5 of the top 20 pathways were related to platelet activation or degranulation (**Extended data table 2**). These included pathways for platelet degranulation, platelet activation, and platelet alpha granules. Additionally, granulocyte migration, neutrophil chemotaxis, neutrophil degranulation, and autophagy were also significantly enriched (**Figure 1D**). This together indicates there is an unique pathophysiology associated with the renal disease in patients with RG blooms.

To further support this distinct RG-driven platelet activation endotype of LN, in contrast, LN patients without RG blooms had over-representation of genes otherwise considered part of canonical pathways of LN disease pathogenesis, including B cell activation, humoral immune response, and complement activation. Evaluating the individual genes that drive related functional pathways, in LN patients with RG blooms, we observed significant upregulation of genes known to play key roles in platelet activation (e.g., alpha-synuclein (SNCA), platelet derived growth factor A (PDGFA), platelet factor 4 (PF4), protein S (PROS1); **Figure 1E, F**) compared to those without detectable dysbiosis. Further, genes related to neutrophil degranulation, but not type-I interferon (**Figure 1H, I**), were also significantly increased in the LN patients with RG blooms. Platelet activation and neutrophil degranulation modules, which define the overall contribution of specific biological pathways in each sample within a gene expression dataset, were also increased in platelet activation and neutrophil module genes in the LN patients with RG blooms, compared to those without gut dysbiosis (**Figure 1G, J**).

Conversely, in the LN patients without blooms, there were instead significant elevations of genes associated with B-cell activation or increased plasma cell differentiation, including MZB1, JCHAIN, and others (**Figure 1E**). Next, to consider whether longitudinal platelet activation changes correlate with LN disease activity, we assessed the platelet activation gene module versus a gene-set module of B cell-related genes. We observed significantly higher platelet activation uniquely in LN patients with RG blooms that correlated with increasing disease activity and was inversely correlated with B cell module genes with disease activity in these same patients (Spearman r = -0.79, p = 0.03, **Extended data 2A**). The platelet-related gene module representation had a significant direct correlation with neutrophil activation genes (r=0.81, p=0.03, **Extended data 2B**).

Interestingly, in a meta-analysis of whole-blood transcriptomics data from patients with active LN (total n=421 patients, with publicly available databases; GSE99967^19^, GSE112087^20^, GSE49454^21^, and GSE65391^22^, we found an inverse relationship between platelet activation/neutrophil degranulation genes and IFN genes (**Extended data 3A**), as well as an inverse correlation between platelet activation/neutrophil degranulation genes and those related to B cell activity. In our studies, this pattern was more pronounced in LN patients with RG blooms compared to those without (**Extended Data 3B, C**). This meta-analysis supported the hypothesis that RG-driven platelet activation represents a distinct pathophysiology of LN disease flares, which validation studies showed was generalizable beyond our initial pilot studies (see below).

### Serum IgG anti-LG antibody is a surrogate marker of gut dysbiosis with RG blooms

Since we were intrigued by the findings in our relatively small pilot cohort of patients and our meta-analysis, we sought to test further the hypothesis that RG blooms in patients with LN drive platelet activation, representing a distinct pathophysiology compared to LN patients without RG blooms. We utilized our previously reported assay^7^, which demonstrated that LN patients with RG intestinal blooms, identified through fecal microbiota analysis, exhibit a robust host systemic immune response characterized by serum IgG2 antibody responses against RG, detectable at dilutions as high as one to one million in sera^1,7^. This serum antibody response is specifically directed against a novel surface lipoglycan made by some RG strains, which is distinct and unique from a large survey of other bacterial glycans^4^, and the serum anti-lipoglycan (Anti-LG) antibody assessment was shown to provide a robust surrogate marker for gut RG blooms in SLE patients^1,7^.

In the validation cohort of LN patients with active disease from the ACCESS trial^23^, at the time of enrollment, all patients had active LN flares, defined by a urine protein-to-creatinine ratio (Pr/Cr) >1, all patients had biopsy-proven class III or IV LN. We therefore stratified patients based on their anti-LG antibody responses (**Figure 2A**). In these LN patients, their baseline level of anti-LG antibodies was generally maintained during longitudinal measurement, over up to 105 weeks^7^. In contrast, anti-LG levels were maintained below a negative cut-off (59,000 U/mL) in our in-house, bead-based quantitative assay (**Extended data 4**). We found no correlations for IgG anti-LG levels based on demographics, antiphospholipid antibody (APL) positivity, complement levels, dsDNA levels, and urine Pr/Cr ratio. There were also no differences in white blood cell (WBC) count, red blood cell count, platelet count, creatinine clearance, albumin, or total urine protein/creatinine ratio (**Extended data table 3**). Furthermore, there was no difference based on the use of hydroxychloroquine (HCQ), mycophenolate mofetil (MMF), azathioprine (AZA), anticoagulants, or prednisone doses, and no patient received more than 40 mg of daily prednisone at the time of sampling.

**Figure 2.**
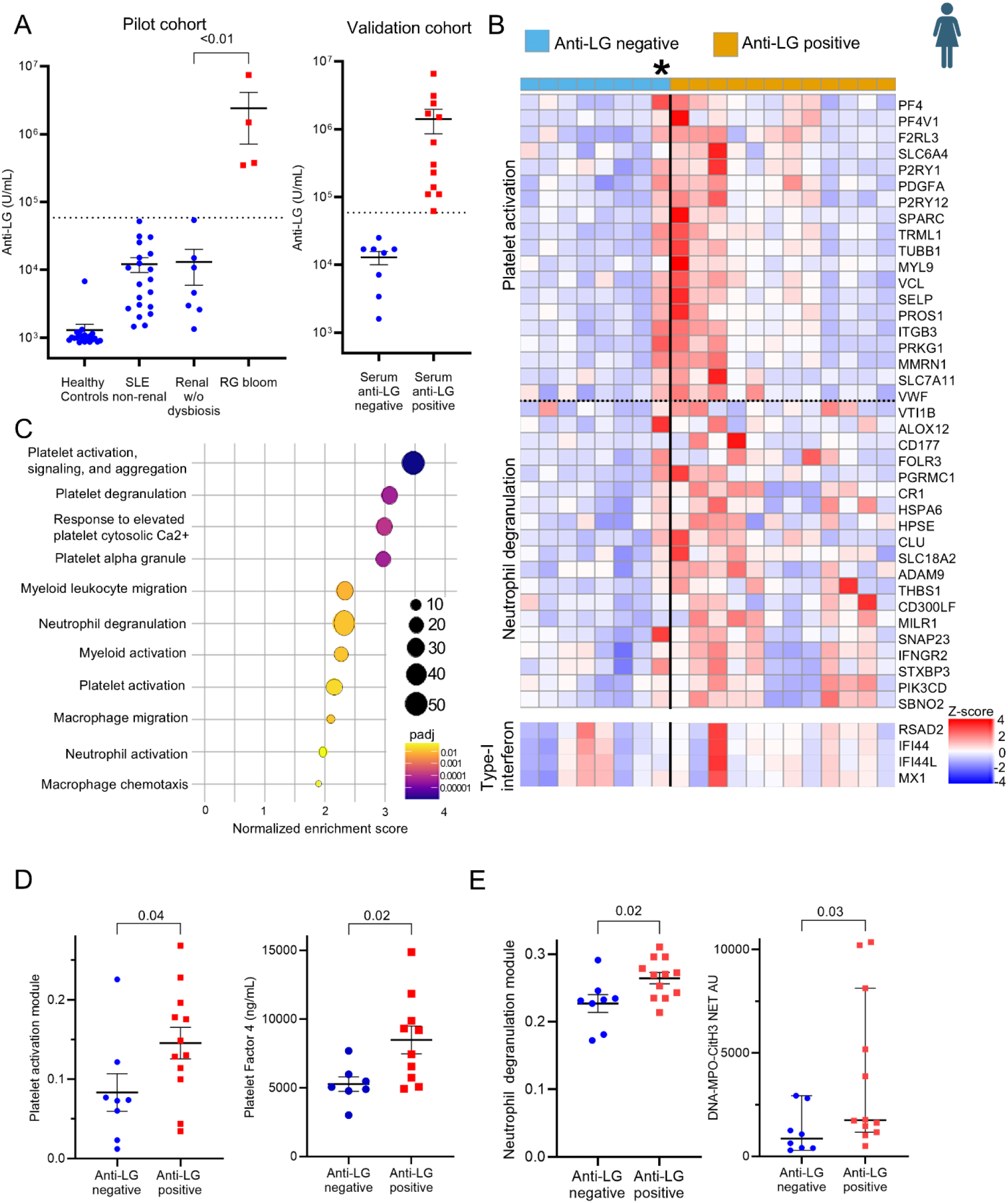
Validation cohort demonstrates platelet activation in patients with anti-LG positive antibody levels. (A) *R.gnavus* (RG) serum anti-lipoglycan (anti-LG) levels in non-renal SLE, patients with LN patients without gut dysbiosis (Renal without dysbiosis), and LN patients with RG gut blooms (RG bloom) from the New York University pilot cohort (NYU)^1,7^. A cut point was used to dichotomize patients based on anti-LG levels (horizontal dashed line) in the ITN034AI clinical trial (ITN) validation cohort, separating those with negative (< 5.9 x 10^4 U/mL) and positive (≥ 5.9 x 10^4 U/mL) levels. (B, C) Genes (B) and pathways (C) related to platelet activation and neutrophil activation increased in anti-LG positive patients. Platelet activation, signaling, and degranulation (NES=3.5, padj=1.02E-06), platelet degranulation (NES=3.07, padj=4.5E-5), response to elevated platelet cytosolic Ca2+ (NES=2.99, padj=1.08E-4), platelet alpha granule (NES=2.96, padj=4.76E-05), myeloid leukocyte migration (NES=2.34, padj=8.08E-03), neutrophil degranulation (NES=2.33, padj=1.22E-02), myeloid activation (NES=2.27, padj=1.25E-02), platelet activation (NES=2.16, padj=2.17E-02), macrophage migration (NES=2.10, padj=1.59E-02), neutrophil activation (NES=1.96, padj=4.97E-02), macrophage chemotaxis (NES=1.9, padj=4.04E-02). *indicates patient with negative anti-LG antibody level based on designation (<5.9x10^4 U/mL), but with elevated urine protein/creatine ratios of >20 g/g. (See Extended data 5) (D) Platelet gene expression module scores (left) and platelet factor 4 (PF4) levels (right) in validation cohort anti-LG negative and positive patients. PF4 levels are shown for patients with class III and/or IV nephritis, without class V disease. (E) Neutrophil degranulation gene module (left) and serum NET fragment detection by ELISA for citrullinated Histone 3 (citH3), myeloperoxidase (MPO), and DNA (right), which are markers for NETosis, in pilot and validation cohort patients with available serum samples. Unpaired, non-parametric t-test.

### In a validation cohort, raised anti-LG antibody levels correlated with platelet activation signature

For validation studies, we used baseline samples from the control group in the ACCESS LN trial^23^. At the time of LN flare, we identified patients with positive anti-LG antibodies^7^, which were found to also have significantly higher expression of genes related to platelet activation and neutrophil degranulation (**Figure 2B, C**). By unsupervised hierarchical cluster analysis, anti-LG positive patients were found to exhibit similar patterns of platelet activation and neutrophil degranulation genes (**Extended data 5A**), which was independent of the expression level of interferon (IFN) stimulated genes (**Extended data 5C**). Interestingly, we identified one seemingly discordant patient with a negative anti-LG antibody level who also exhibited a high platelet activation signature in the whole blood RNAseq libraries. Strikingly, this patient had a massively abnormally elevated urine protein-to-creatinine ratio (i.e., >20 g/g) (**Figure 2B, Extended data 5B**), which may suggest that urinary loss interfered with the detection of this serum anti-bacterial response.

Whole blood RNAseq data were available for the control group of patients with class III and class IV glomerular disease at only the time of enrollment. In this validation cohort in LN flare, we observed a subset of patients with the same platelet gene module identified in our test cohort, and these patients also had significantly higher serum platelet factor 4 (PF4) levels in the patients with raised serum anti-LG positive antibodies, compared to the anti-LG negative patients (**Figure 2D, Extended data 5D, Extended data 6**). Notably, the release of PF4 from activated platelets can play a critical role in downstream myeloid cell activation^24^.

Next, we directly considered how RG-driven platelet activation may lead to changes in granulocyte effector functions^24^. We found that LN patients with RG bloom also had raised serum anti-LG antibodies (**Figure 1E, left**). In whole-blood RNA sequence libraries, these patients had gene signatures reflective of increased neutrophil degranulation, and also had serum levels of the NET fragments, myeloperoxidase (MPO), citrullinated histone H3 (citH3), and dsDNA, which all were significantly higher than those found in anti-LG negative patients (**Figure 1E, right; Extended data 5E, Extended data 6**).

We then considered which of the elevated gene transcripts may better correlate with the elevated levels of serum proteins that also provided evidence of platelet and/or neutrophil activation. Significant correlations were found between PROS-1 (r=0.52, p=0.02), PPBP (r=0.48, p=0.03), and TREML1 transcript (r=0.48, p=0.03) with serum PF4 protein. There was also a significant correlation of EGF (r=0.53, p=0.04) and KRT1 transcripts (r=0.58, p=0.02) with serum protein detection of NET fragments (**Extended data 7**). These findings demonstrated a direct relationship between platelet activation-related transcripts and protein-level evidence of platelet and neutrophil activation.

### Intestinal RG colonization induces platelet activation

To assess potential causal relationships and determine whether RG strains from patients with LN can directly affect platelet activation, we performed *in vivo* challenge studies in mice, applying methods from our earlier report^11,25^. After oral antibiotic preconditioning that facilitates intestinal colonization (**Figure 3A**), at 2 weeks after gavage treatment, we documented higher levels of fecal RG in the mice that received oral RG strains from patients with LN, but not in mice that received sham gavage, we documented higher levels of fecal RG by species-specific qPCR (**Figure 3B**). Notably, two weeks after oral RG gavage, increased gut permeability was observed following RG colonization (**Figure 3C**). This was indicated by higher serum levels of FITC-dextran three hours post-introduction into the gut, which demonstrated that RG induced increased gut permeability. This effect was further confirmed by correlated increases in the serum levels of a gut microbial product, LPS-binding protein, which reflects heightened intestinal permeability^26^ (**Extended data 8**). At this time point of high RG colonization, we performed whole blood RNA sequencing. Principal component analysis (PCA) of the top 25% most variably expressed genes showed distinct gene expression profiles in the whole blood of mice with RG gavage compared to those with sham gavage (**Figure 3D**). Like patients with LN and RG blooms (**Figure 1&2**), mice after RG gavage also demonstrated significantly increased expression in gene transcripts reflecting platelet activation (**Figure 3E, F**).

**Figure 3.**
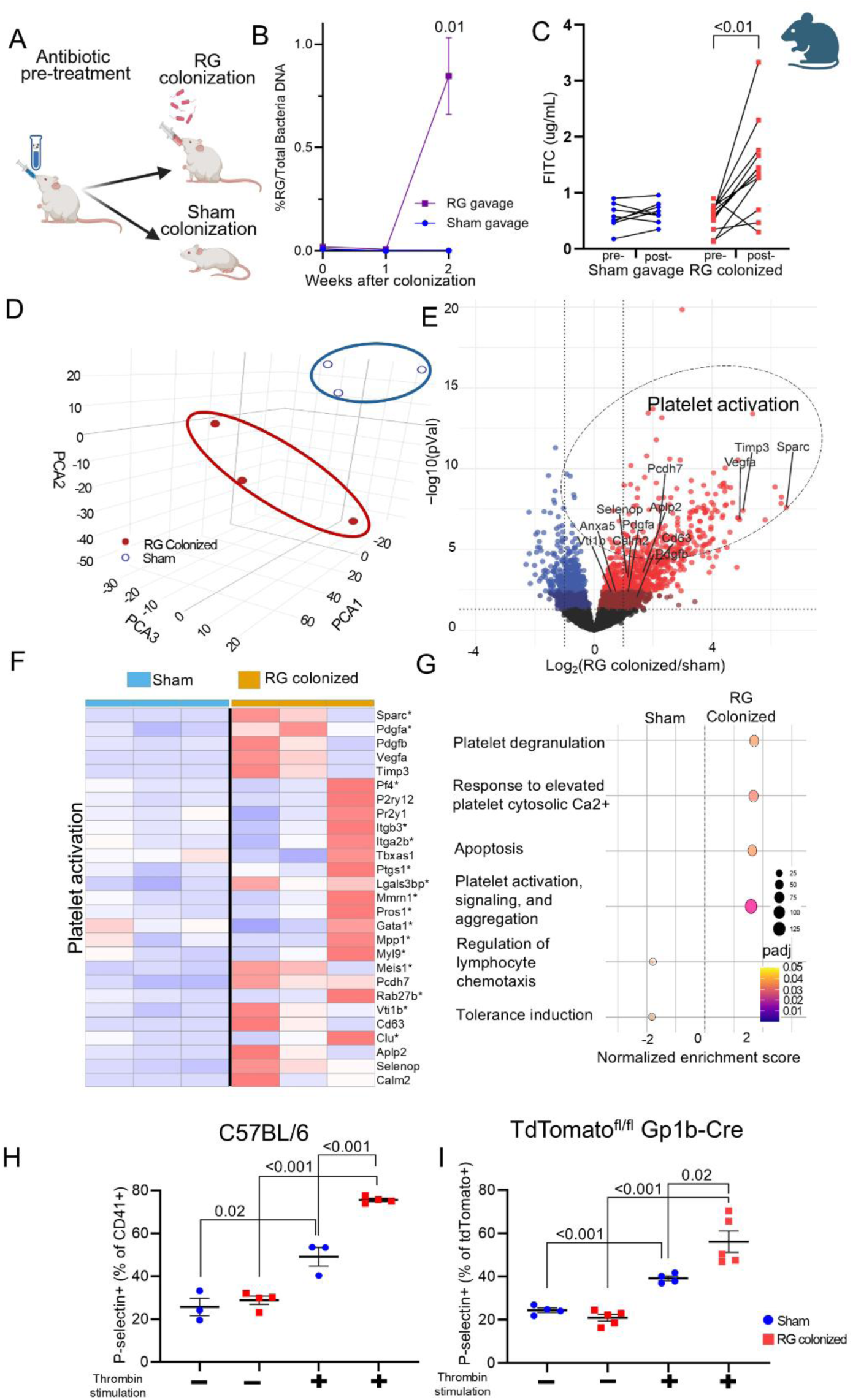
Flow cytometry and immunohistochemistry show functional changes in platelet activation after RG gavage. (A) Experimental model of antibiotic pretreatment, subsequent gavage of RG or sham, and sample collection, (B) Assessment by PCR of stool *R. gnavus* throughout the experiment, identifying peak colonization of RG at 2 weeks after the last gavage in C57-BL/6 mice. Mice were pre-treated for two weeks with an antibiotic cocktail of enrofloxacin, ampicillin, and metronidazole. Mice were then gavaged with either sham or RG every other day, for a total of 5 administrations over two weeks. Timepoint 0 represents the end of this gavage period. (C) FITC-dextran levels in serum pre- and post-oral gavage of sham vs. RG gavage-treated (RG colonized) mice. (D) Principal component analysis of the top 25% varied genes among C57-BL/6 mice 2 weeks and 4 weeks post-gavage with either sham or RG. PCA1 48.7%, PCA2 28.3% and PCA3 12.1%. (E) Gene expression 2 weeks post-gavage with genes related to platelet activation noted. (F) Heatmap of whole-blood RNAseq genes related to platelet activation in sham gavage (left) and RG colonized (right) mice. *indicates transcripts identified as significantly increased in murine challenge studies, as well as human whole-blood RNAseq studies (Figure 1,2). (G) GSEA pathway analysis of genes overexpressed in RG compared to sham at 2 weeks post-gavage. (H, I) P-selectin+ platelets (relative to CD41+ FSC ^low^ in C57-BL/6 mice (H) and Gp1b-Cre+ platelets in TdTomato-Gp1bCre mice (I) with or without Thrombin stimulation. Unpaired t-test.

By GSEA pathway analysis, genes related to platelet activation and degranulation were significantly overrepresented in mice that received RG gavage compared to mice that received sham gavage (**Figure 4G**; platelet degranulation, NES=1.7, padj=2.4E-2; platelet activation, NES=1.6, padj=6.5E-3). No differences were found in the RG-colonized vs. sham-gavaged mice in WBC count, lymphocytes, neutrophils, hemoglobin, platelets, or in the mean platelet volume were observed (**Extended data 9**).

**Figure 4.**
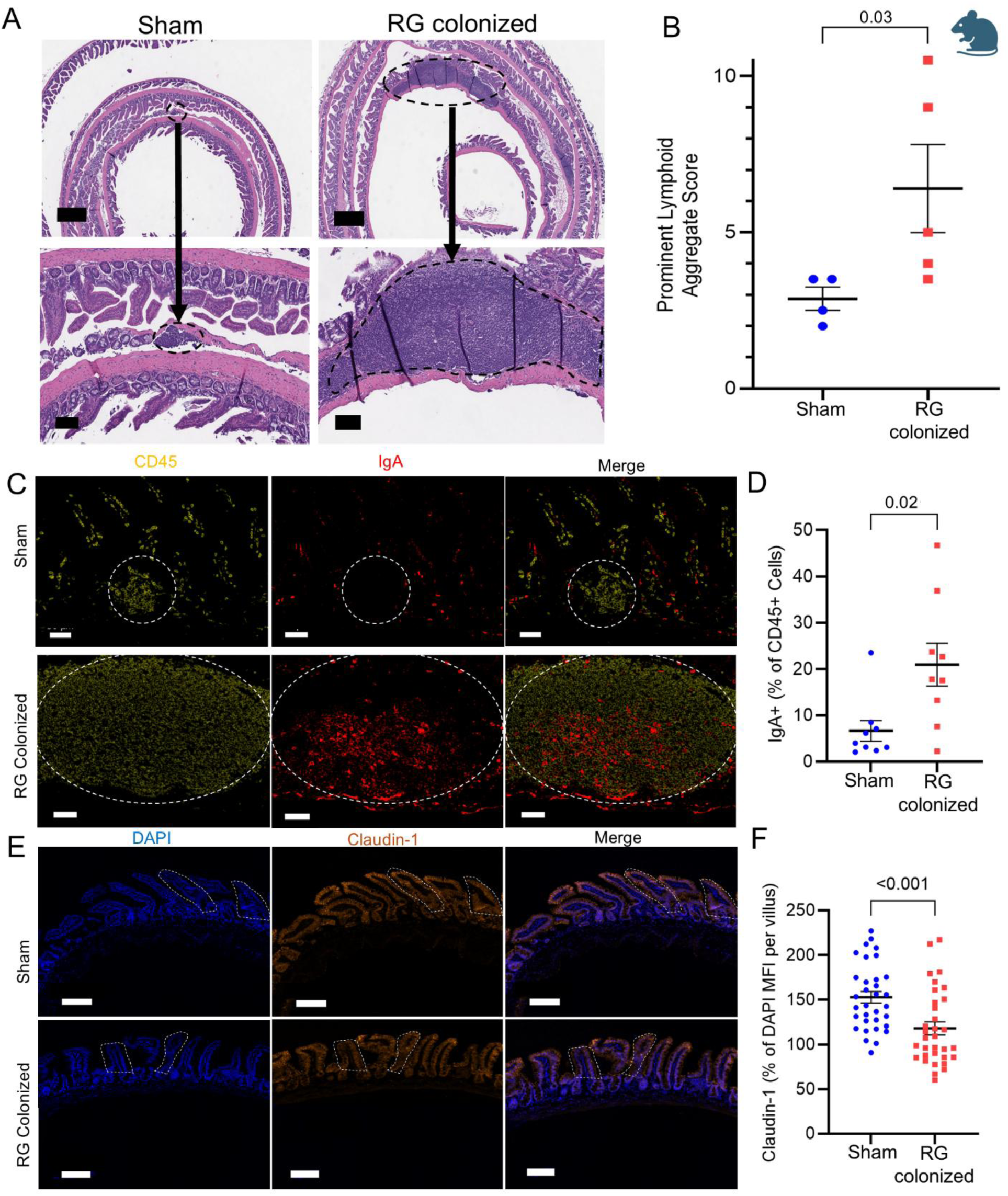
*R. gnavus* gavage drives a robust immune response *in vivo* challenge studies. (A) Lymphoid aggregates in the terminal ileum in sham (left) vs. RG-colonized (right) mice at the terminal ileum. Bar = 50µm top, 100µm bottom. (B) Quantification of total lymphoid aggregates in Swiss rolls, with scoring weight (range 2+ to 10.5+) determined by number of lymphoid cells per aggregate. (C, D) Immunofluorescence of relative IgA level, as a ratio of total CD45+ lymphocytes per aggregate, bar = 50µm and (E, F) of MFI of Claudin-1 intensity relative to DAPI MFI per villus as a measure of relative tight junction quantification. Bar = 100µm. Non-parametric, unpaired, t test.

To begin to consider elucidating how RG-driven platelet activation may contribute to disease progression, we investigated the responsiveness of RG-primed platelets to *ex vivo* challenge various agonists. By flow cytometry, we quantified levels of surface P-selectin, a well-established marker of platelet activation (**Extended data 10**). In both C57BL/6J mice and transgenic mice with platelets marked by tdTomato-Gp1b, following RG gavage there was significantly higher surface levels of surface P-selectin expression in mice compared to sham-treated mice after platelets were stimulated with human plasma thrombin, a potent activator of PAR1 and PAR4 receptors (**Figure 3H, I**). No differences in platelet activation levels were observed with PBS. In contrast, these platelets did not show functional differences after subsequent challenge with equine type I collagen fibrils that primarily activate GPVI, nor with ADP that signals through P2Y1 and P2Y12 receptors (**Extended data 11**). Hence, RG activation may involve platelet-leukocyte interactions and thrombus formation that is specific to thrombin-PAR1/PAR4 pathway, as opposed to effects in platelets affected through GPVI (Collagen stimulated) or P2Y1/P2Y12 (ADP stimulated) pathways.

### RG colonization leads to a robust intestinal immune response

To consider the mechanistic link between gut colonization with RG and evidence of increased peripheral platelet activation, we examined the terminal ileum in the experimental mice and documented that RG gut colonization induced the development of prominent large lymphoid subepithelial follicles in the distal small intestine (**Figure 4D, E**). These lymphoid structures in LG-producing RG-colonized mice had an increased representation of IgA+ B-lineage cells, likely plasma cells (**Figure 4F**). Between epithelial cells in the villi lining the gut, lower expression levels of the tight junction protein, claudin-1, were also observed in the RG-colonized mice (**Figure 4H**), rationalizing the impairment of intestinal barrier functionality that followed intestinal colonization with LG-producing RG strains.

### Intestinal colonization leads to increased megakaryocytes

Given that platelets have a half-life of 7-10 days in humans and somewhat shorter in mice^27^, they still showed notable functional differences when responding to thrombin stimulation two weeks after the last RG gavage exposure. We hypothesized that RG colonization might affect the turnover of platelet cellular precursors, namely megakaryocytes, which may account for these functional alterations. Further, changes in megakaryocytes following RG exposure are particularly significant when considering long-term platelet effects on the immune system. To explore this possibility, we examined the spleens of mice treated with RG gavage for changes in megakaryocyte numbers, as the spleen is a key site of extramedullary hematopoiesis and platelet production. We observed a significant increase in the number of megakaryocytes in both the bone marrow (**Figure 5A**) and the spleens of mice treated with RG gavage compared to those with sham gavage (**Figure 5B**). We next assessed lupus-prone FcγRIIb^-/-^ congenic mice^28^ that were either sham-treated or colonized with RG strains, including isogenic paired strains that differ only in LG production. We also observed increased megakaryocyte numbers in the spleens of mice colonized with LG-producing pathogenic strains of RG, but not in mice colonized with a RG strain (nb90) that does not produce the LG toxin (**Figure 5C**).

**Figure 5.**
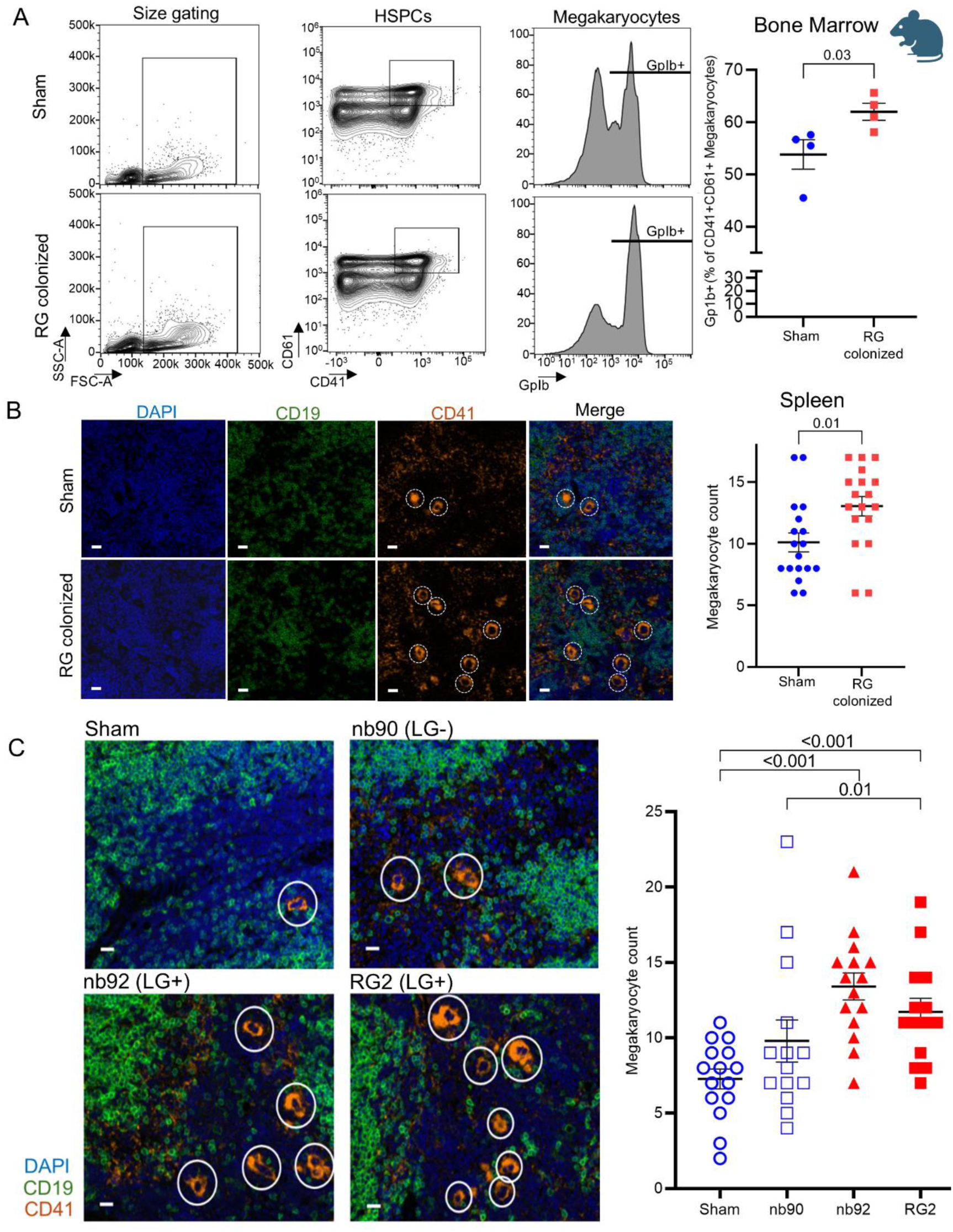
Increased megakaryocytosis with RG-colonization. (A) Flow cytometry of bone marrow megakaryocytes in TdTomato-Gp1bCre mice at 2 weeks post-sham vs. RG oral gavage. (B) Immunofluorescence of megakaryocyte numbers in 3-4 months old, female TdTomato-Gp1bCre mice after RG gavage as defined by CD41+. (C) Immunofluorescence of splenic megakaryocyte in 12 month old, female, FcγRIIb^-/-^ mice as defined by CD41+ after gavage with LG-positive RG strains (nb92, RG2) versus strains without LG (nb90). (B,C) Cell counts represent counts per high-powered field, at least three mice per group. Unpaired, non-parametric t-test.

### Pathogenic LG-producing RG strains induce renal cellular infiltration

Our previous study^11^ investigated how RG colonization affects lupus progression in the B6.Sle1.Sle2.Sle3 triple congenic (TC) mouse model, which develops severe, fully penetrant systemic autoimmunity and glomerulonephritis driven by Sle1 (loss of nuclear antigen tolerance), Sle2 (lowered B cell activation), and Sle3 (CD4+ T cell dysregulation and anti-dsDNA antibody production). However, since these mice spontaneously develop severe renal disease, we sought to test the impact of gut colonization in a lupus-prone model that does not otherwise develop significant renal pathology. Indeed, FcγRIIb^-/-^ is the only Fcγ receptor on B cells in both mice and humans, and its critical role as an inhibitory receptor is highly conserved. Its absence in mice leads to unrestrained activation of autoreactive B cells by IgG immune complexes, effectively accelerating systemic autoimmunity. The relevance of this model to human lupus is further underscored by genetic studies identifying polymorphisms in human FcγRIIb^-/-^ that increased susceptibility to SLE^29^. We therefore evaluated the role of RG in the genetically susceptible, well-validated FcγRIIb^-/-^ murine model ^28^, which does not develop severe renal disease without an additional insult.

Mice colonized with LG-producing RG strains showed worsening renal pathology as demonstrated by increased tubulointerstitial mononuclear cell infiltrates in mice that had been gut colonized with LG-producing RG, but not by non-producing strains (**Figure 6)**. These mice also exhibited increased evidence of neutrophil degranulation (NETosis), as indicated by the deposition of myeloperoxidase and citrullinated histone 3. Increased renal macrophage cell and T cell infiltration was also observed by immunofluorescence (**Figure 7**).

**Figure 6.**
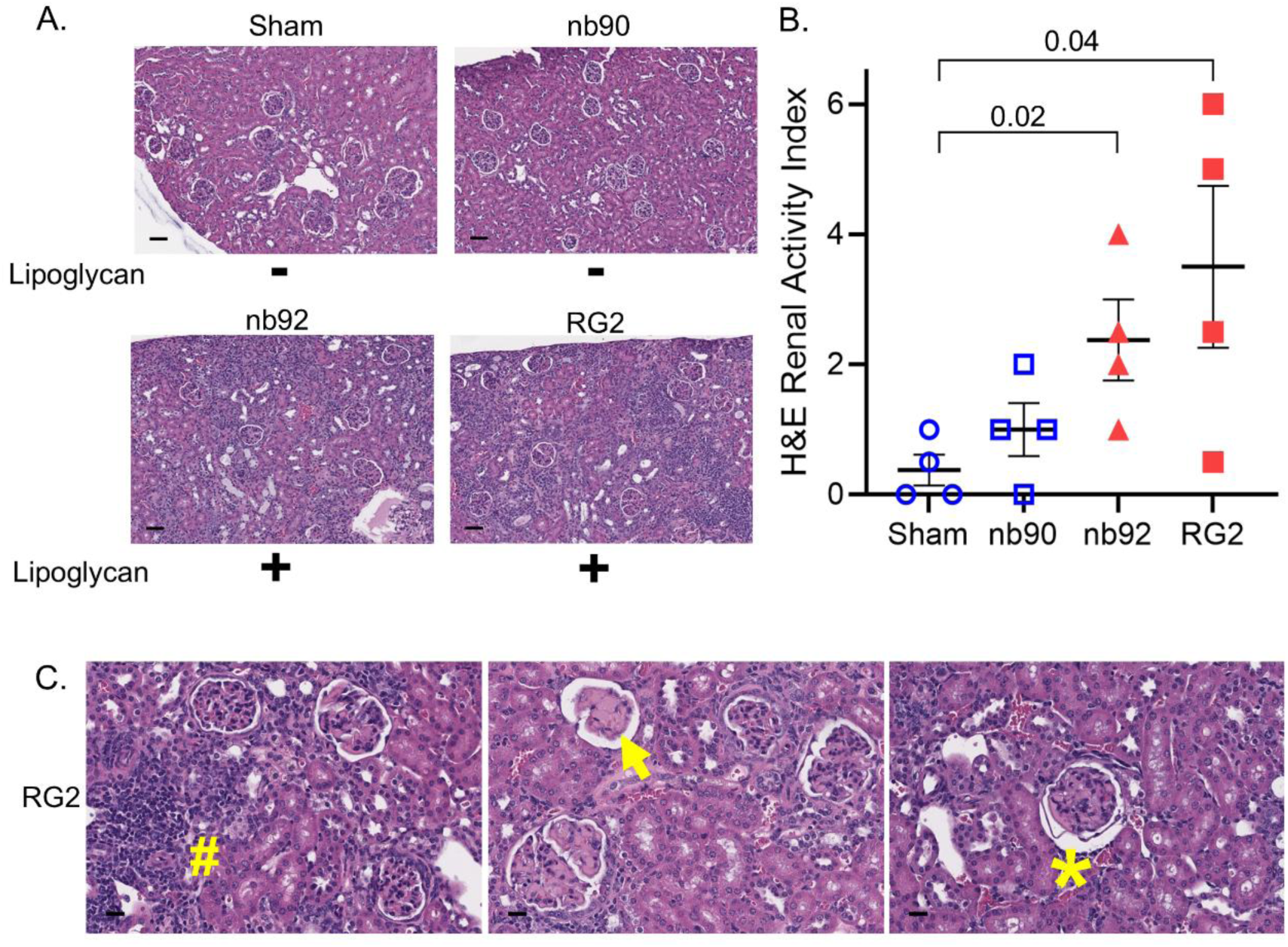
*R. gnavus* gut colonization causes increased renal cellular infiltration in lupus-prone mice. (A) Representative kidney sections (H&E) after FcγRIIb-/- colonization with lipoglycan-positive *R. gnavus* strains (nb92, RG2), and lipoglycan-non-producing isogenic (nb90) strains, ten days after the end of colonization. (B) Quantification of the renal activity index is scored based on a blinded review for the presence (scored from 0-3) of mononuclear cells in the tubulointerstitium, glomerular leukocyte infiltration (#), and deposition of hyaline thrombi (Arrow), closed capillary loops, or early crescent formation/fibrotic ring (*). (C) 40x magnification example glomeruli of RG2 (LG-producing RG strain) colonized FcγRIIB-/- lupus-prone female mice, showing hyalinization (left, middle), and mononuclear interstitial cellular infiltrations (right).

**Figure 7.**
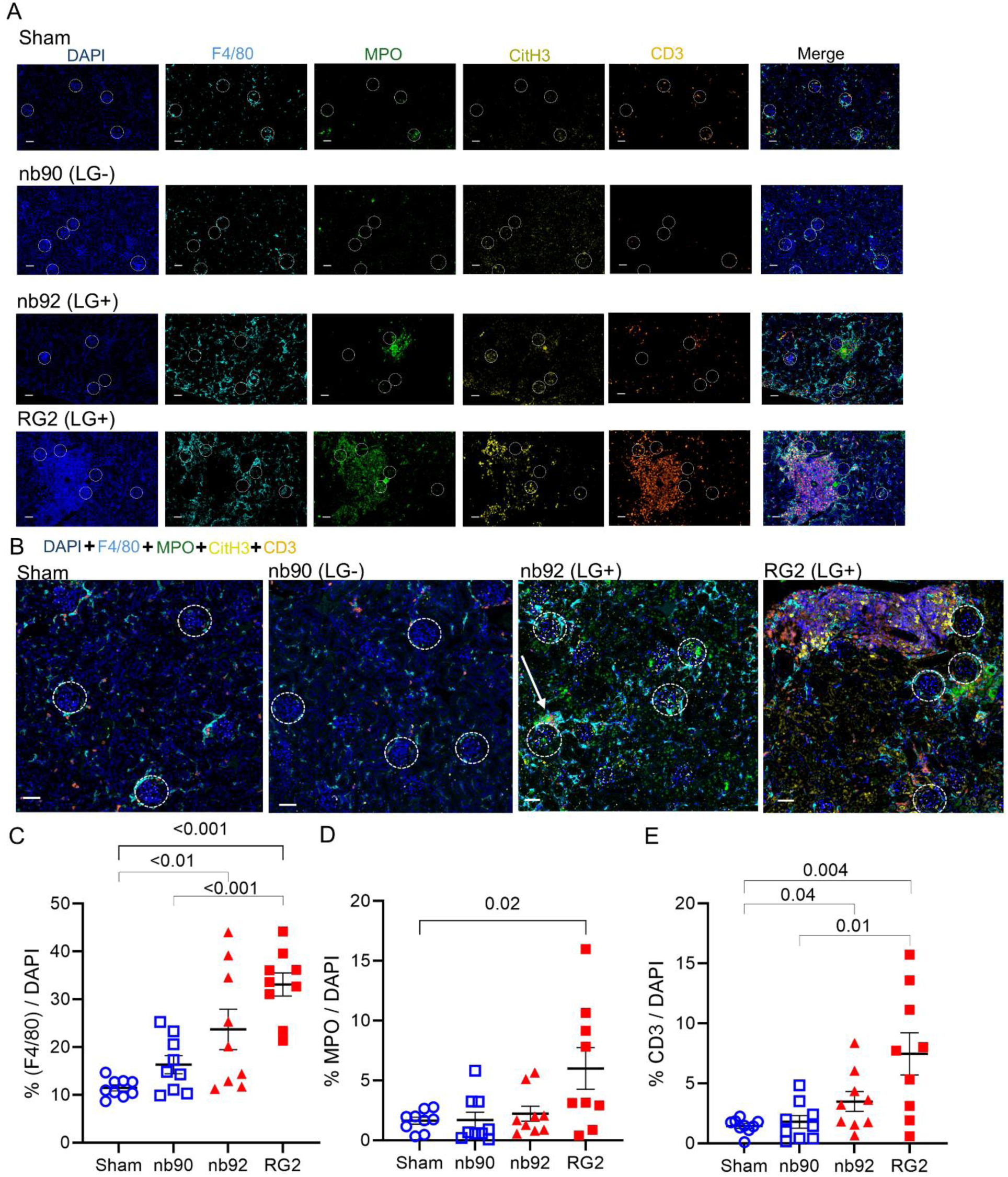
*R. gnavus* gut colonization causes increased renal-resident monocytes and increased evidence of NETosis. (A) Representative imaging for DAPI (all cells), F4-80 (macrophages), MPO (neutrophils), citrullinated H3 (neutrophils-NETosis), and CD3 (T cells) after FcγRIIb-/- colonization with lipoglycan-positive *R. gnavus* strains (nb92, RG2), and lipoglycan-non-producing isogenic (nb90) strains. (B) Quantification of cell percentage of F4-80 (macrophages)+ cells relative to all cells, (C) Cell percentage of MPO+ (neutrophils) relative to all cells. (D) Merged immunofluorescence of macrophages (F4-80), neutrophils (MPO), with evidence of NETOsis (CitH3), and T cells (CD3). Arrow highlights area of NETosis adjacent to glomeruli, based on MPO, CitH3, and F4-80 detection.

### The LG produced by RG itself drives *in vivo* platelet activation and megakaryocytosis

To determine whether the RG surface expressed LG toxin itself is sufficient to drive changes *in vivo*, specifically for splenic megakaryocytosis, we administered purified LG intraperitoneally in a well-validated lupus-prone mouse model (FcγRIIb^-/-^ mice^28^), and assessed the impact on megakaryocytosis 6 days after intraperitoneal challenge with a LG preparation from the RG2 strain. Akin to C57BL/6J mice after RG gavage, we also observed that injection of LG also drove increased platelet responsiveness to thrombin stimulation (**Figure 8A, B).** In addition, a marked level of megakaryocytosis (i.e., expansion of the representation of the cellular precursors of platelets) was detected in the spleens of these RG-challenged lupus-prone mice, which was not detected after control treatment (**Figure 8C**). Compared to intraperitoneal injection with vehicle alone, LG-injected mice exhibited neither differences in WBC count, lymphocytes, neutrophils, hemoglobin, platelets, nor mean platelet volume **(Extended data 12**).

**Figure 8.**
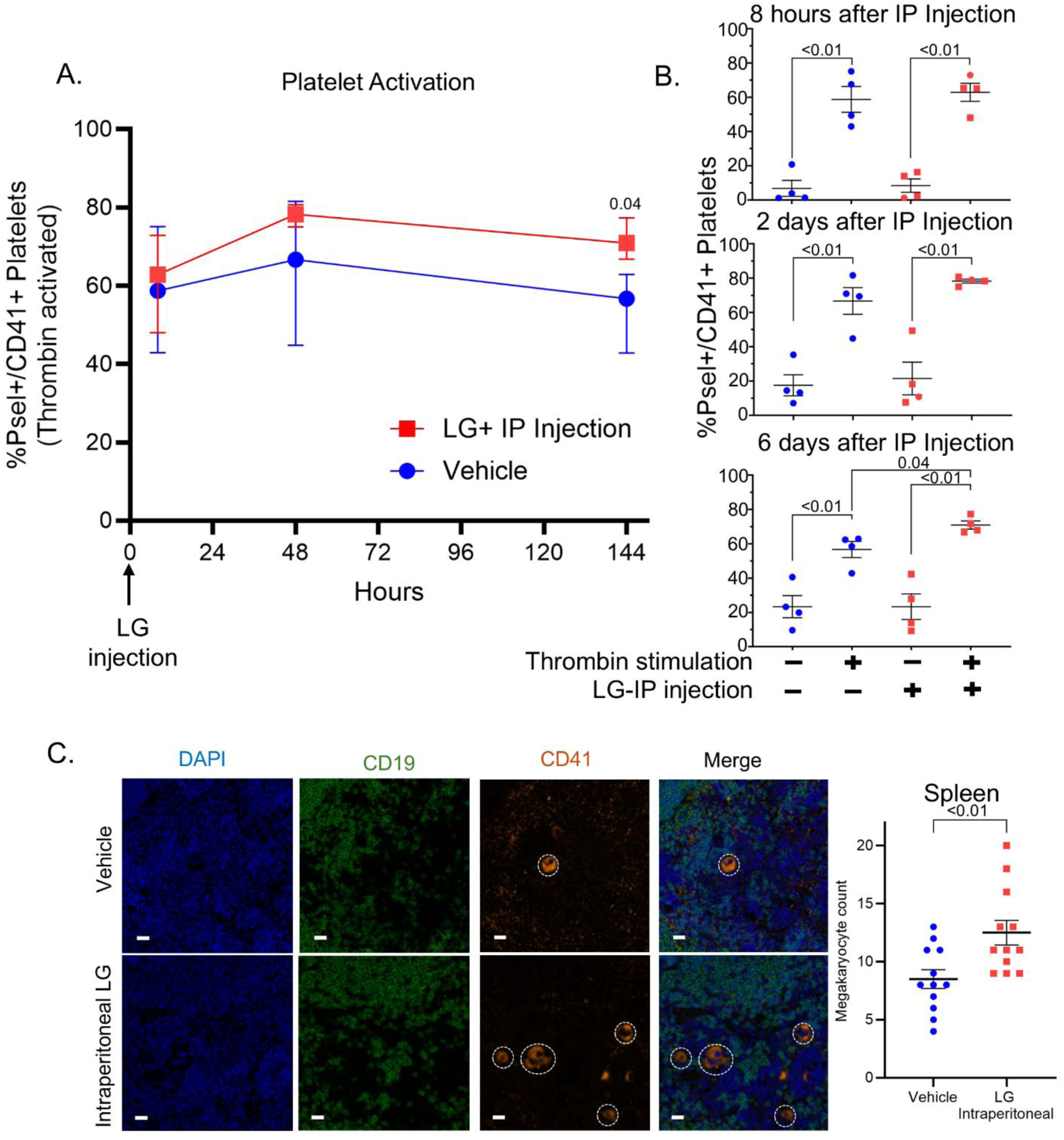
Intraperitoneal RG LG administration increases megakaryocyte numbers in the spleen. (A,B) Platelet activation induced by a single intraperitoneal injection of lipoglycan (1.5µg/g) in FcγRIIb-/- mice was evaluated by flow cytometric monitoring of the percentage of surface P-selectin expressing cells after thrombin stimulation at 8h, 2 days (48h), and 6 days (144h). (A) Shows mean with range. IP: intraperitoneal. LG: lipoglycan. n=4. (C) IF megakaryocyte quantification in the spleen of 5-8month old lupus-prone, female, FcγRIIB-/- mice six days post one-time LG intraperitoneal injection (2µg/g). Unpaired, non-parametric, t-test.

Cumulatively, our results show that, akin to the events in a major subset of LN patients, LG-producing RG blooms in the murine gut induce increased platelet activation. The transcriptomic profiles induced by this RG colonization are also akin to those in RG-exposed patients, as transcriptomic profiles are distinct and distinguishable from the canonical expression pathways observed in LN patients without RG blooms. **Figure 9** provides a graphical summary of the mechanistic model assessed by these experiments, illustrating how RG specifically drives platelet activation and can subsequently contribute to LN disease pathogenesis in a subset of patients with SLE.

**Figure 9.**
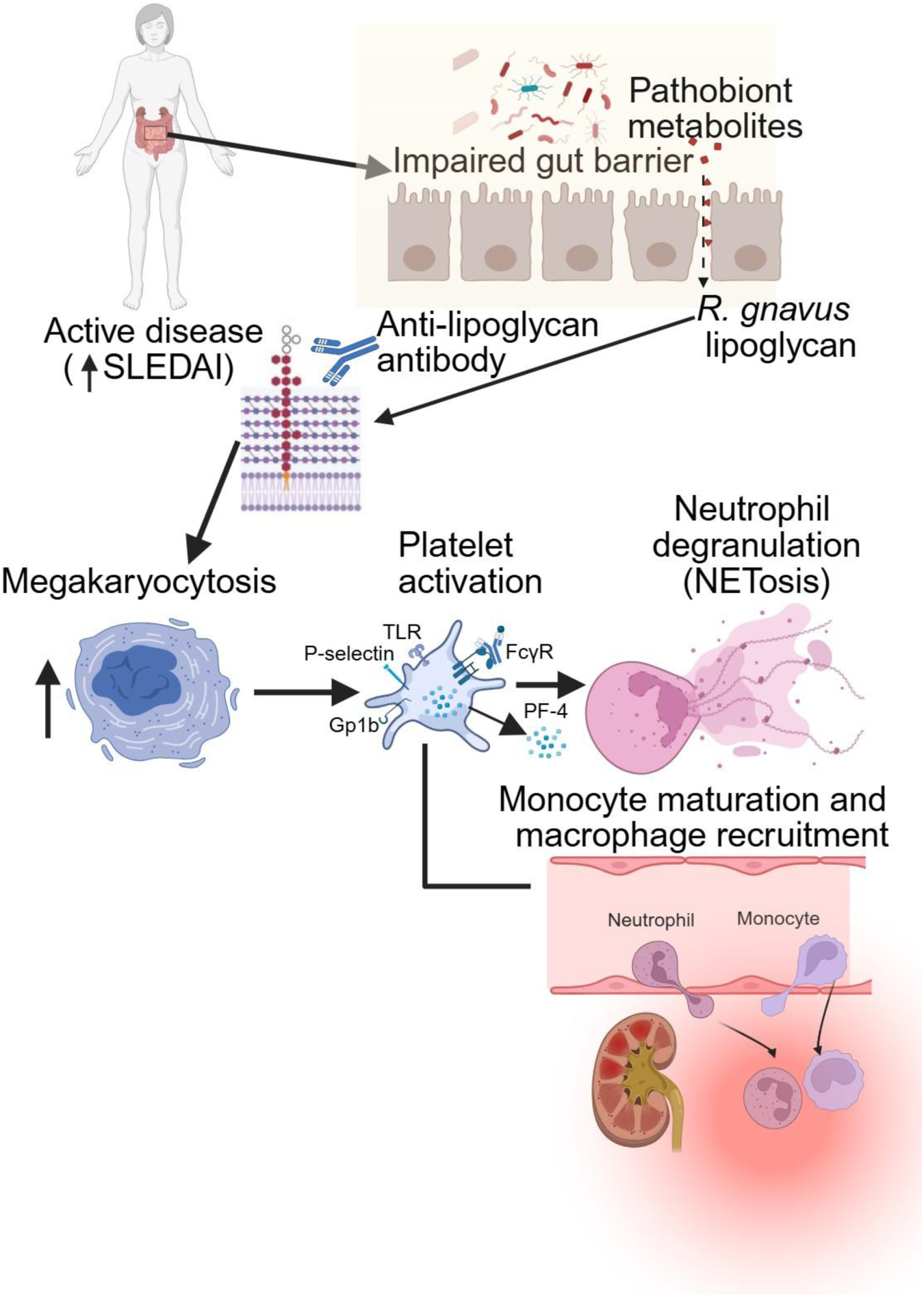
A model of the mechanistic hypothesis in SLE. Impaired gut barrier function allows microbial products to translocate, exposing the immune system to RG surface lipoglycan (LG). RG can activate platelets via TLRs and bind FcγR, promoting platelet-immune cell interaction. This leads to LN damage through platelet-derived extracellular vesicles priming neutrophils to release neutrophil extracellular traps (NETosis). Activated platelets also release soluble factors such as platelet-factor 4, aiding granulocyte maturation and sustaining inflammation. Abbreviations: sP-selectin, soluble P-selectin; RG, *Ruminococcus gnavus;* SLE, systemic lupus erythematosus; LN, lupus nephritis; SLEDAI; Systemic Lupus Erythematosus Disease Activity Index; NET; Neutrophil extracellular traps. Adapted from^4,24^.

## Discussion

In this study, we identified distinct pathophysiological features in lupus nephritis (LN) patients with gut dysbiosis due to expansions of the microbial commensal *Ruminococcus gnavus* (RG). Our unbiased survey of whole-blood RNA sequencing libraries from patients from our prior longitudinal study^7^, revealed distinct transcriptomic signatures during peak intestinal expansions of RG. Herein, we studied RNA isolated directly from whole blood without fractionation, which provided a comprehensive view of the immune landscape in each LN patient. LN patients with RG blooms exhibited a distinct gene expression profile, characterized by the activation of platelet pathways and neutrophil degranulation. In contrast, LN patients without RG blooms instead displayed canonical gene expression patterns of B cell and complement activation (**Figure 1**), which have been generally highlighted in earlier reports^20,22^. Yet, in publicly available whole-blood gene expression datasets with active LN, with new criteria from our pilot and validation cohorts, we identified patients with the same features of this newly identified endotype, and this included relatively heightened platelet and neutrophil activation, along with relatively lower interferon and B-cell activation gene signatures.

The utility of the anti-LG antibody assay as a surrogate marker for host immune exposure to intestinal RG blooms was also documented in both our pilot and validation cohorts. In a major LN subset within this cohort, anti-LG antibody levels were consistently elevated over time, correlating with increased disease activity and platelet activation. This marker, therefore, provided a means for identifying LN patients with RG-driven platelet activation, without the need for 16s-rRNA sequencing in our validation cohort. Positive anti-LG levels and platelet activation were highly correlated, demonstrating persistence of high anti-LG antibody levels at both cross-sectional and longitudinal measurements extending up to two years. These measurements proved to be reliable predictors of RG-driven platelet activation, as measured by whole-blood RNA sequencing, in LN patients with class III or IV disease who had less than 40 mg of daily prednisone exposure. The generalizability of the anti-LG antibody to stratify distinct endotypes of LN, based on inversely related platelet/neutrophil gene versus interferon/B cell signatures, is also supported by the meta-analysis of whole-blood expression data from ^19^ ^22^, which emphasized the detection of neutrophil^19,20,22^ and/or interferon signatures^20–22^ in subsets of patients with LN. Platelet-related gene sets were mentioned in Figgett et al. 2019^20^, but not as key pathophysiological mechanisms of LN disease flares. Additionally, these previous studies did not offer a serum biomarker to differentiate one subset from another, while we have developed and validated the anti-LG antibody assay may provide a straightforward serum blood test for the stratification of pathologic LN endotypes.

In our studies, we focused on LN patients with proteinuria exceeding 1 g/g at the time of sample collection, an indicator of active LN. Notably, one patient exhibited nephrotic syndrome with a urine protein to creatinine ratio over 20 g/g, had a negative serum anti-LG lipoglycan level, but nonetheless showed evidence of marked platelet activation, and associated downstream effects on neutrophils. This underscores the importance of considering additional factors when assessing LN individuals. We postulate that this could reflect antibody loss due to proteinuria from the renal disease. This atypical patient presentation emphasizes the need for further evaluation in the context of high proteinuria regarding other factors, such as the potential concurrent loss of anti-thrombotic proteins. Our findings highlight the use of anti-LG antibody assessments to identify LN patients with gut dysbiosis and to guide therapeutic strategies targeting microbial dysbiosis and platelet activation.

Our study investigates the mechanistic pathways through which RG blooms drive platelet activation. In LN patients experiencing RG blooms, we documented heightened expression of neutrophil activation genes. Our transcriptomic data were substantiated by protein-level assessments, with serum analysis confirming the presence of proteins indicative of platelet activation, with PF-4 release, and neutrophil activation, with NETosis markers such as MPO, citrullinated histone H3, and dsDNA. Hence, RG-driven platelet activation was also linked to increased neutrophil degranulation.

Gene expression profiles from both human cohorts and mouse models revealed pathways enriched for platelet activation. Additionally, functional changes in platelet activity, particularly in response to thrombin stimulation, provide *in vivo* mechanistic support for the hypothesis that RG colonization primes platelets to exhibit overt, excessive responses to stimuli. This RG-driven thromboinflammation leads to negative downstream effects, which are also crucial for understanding cardiovascular risk in SLE^12^. RG colonization, along with the introduction of the RG lipoglycan (LG) preparation itself, drives megakaryocytes, underscoring that RG exposure can have a long-term impact on the critical role of platelets in SLE immune system dysregulation. Identifying RG blooms as a distinct pathophysiological feature in LN patients is significant, as it points to an otherwise unknown environmental trigger of LN flares. Whereas activated neutrophils play a crucial role in trapping and eliminating pathogens, dysregulated excessive levels of NET formation can lead to inflammation and tissue damage, which have been implicated in the etiopathogenesis of SLE. While triggers of such pathways have been poorly understood, our findings suggest that gut dysbiosis may be a powerful driver.

We found functional changes in platelet activity, especially in response to thrombin stimulation, that support the hypothesis that RG colonization primes platelets for excessive responses to stimuli, leading to negative downstream effects on neutrophils. Platelets from RG-colonized mice showed higher activation in response to thrombin than sham-treated mice, indicating a specific pathway of hyperactivation. This underscores the RG LG toxin as a key pathogenic factor. Introducing RG LG preparations alone induced platelet activation and megakaryocytosis, demonstrating RG’s long-term influence on platelets in SLE immune system dysregulation.

Overall, the evidence suggests that RG-induced platelet activation may play a significant role in LN pathogenesis, particularly through its downstream effects on neutrophil function, for inducing NETosis and inflammation. In the kidneys of RG-colonized lupus-prone mice, we observed increased neutrophils, as well as macrophages and T cells among infiltrating immune cells.

While we will further assess the characterization of these macrophages (e.g., M1 vs. M2) and consider if the infiltrating CD3 cells have pathogenic features (e.g., Th17-like), we recognize this finding just weeks after colonization with RG as a model of early-stage immune cell infiltration driven by gut dysbiosis. This condition differs from typical glomerulonephritis observed in humans, which progresses over prolonged periods of immune activation, ultimately resulting in renal disease, particularly in proliferative lupus nephritis. Although full disease manifests from a long-term process of accumulating effects over years, repetitive insults with TLR stimulation^7^ (e.g., RG LG) may lead to renal injury typical of classic glomerulonephritis. Ultimately, we must consider the implications in lupus of defects in apoptotic clearance, NETosis, and myeloid activation, initially triggered by RG-driven platelet activation.

Conceptually, the findings presented in this study contribute to the linked-intestinal bloom-related autoimmunity (LIBRA) syndromes framework^4^, which draws parallels with post-streptococcal autoimmune syndromes (e.g., rheumatic fever and post-streptococcal glomerulonephritis). Unlike overt infectious processes, expansions of species in dysbiotic gut microbiota communities in SLE trigger host immune responses by compromising the gut barrier integrity through otherwise commensal bacteria, such as RG in LN. Historically, these gut microbes were generally undetectable in routine clinical exams, allowing pathobiont expansions to persist and develop over months or years. We present an antibody-based assay designed to identify a specific subset of LN patients with dysbiosis and define the mechanistic downstream pathways of immune activation. This assay lays the groundwork for future interventions addressing microbial dysbiosis in LN and the subsequent platelet activation it triggers. Utilizing this marker can help categorize patients by their risk of disease flare-ups and inform therapy decisions, including drugs targeting microbial dysbiosis and platelet activation. Focusing on RG pathways presents a potential strategy for certain LN patients.

This study has several limitations, including small cohort sizes and the use of archived samples. In longitudinal patient studies, we observed *in vivo* effects reflecting prolonged RG immune exposure, and there is now a need to better assess the dynamics of gut expansion and persistence of defined RG strains, along with the need for longitudinal host gene expression data extending beyond the initial flare. The mouse models, including C57BL/6J mice and lupus-prone murine models, provided valuable insights but were of short exposure, which likely explains why glomerular lesions were not detected. Experimental design can be modified to address the consequences of prolonged intestinal RG persistence.

Furthermore, while we know that RG colonization is almost ubiquitous in the human gastrointestinal tract of infants , there are unaddressed questions regarding host-RG immunobiologic interactions, associated with later temporal shifts between RG strains, particularly during disease quiescence versus flares, and the relationship with presumed prolonged exposure to the LG toxin in LN patients, and the persistence or decay of B-cell anti-LG responses. Longitudinal studies will therefore be essential to assess the ecobiology of RG colonization, competition between species, and strain shifts over time in response to alterations in metabolomic factors and other transitions during disease flares.

Overall, our study highlights the unique pathophysiology linked to RG blooms in LN patients. Therefore, these findings bring into focus potential clinical implications of RG-induced pathways that could be targeted, ideally without broader immunosuppressive influence, as is now standard of care for lupus. The presence of RG blooms and associated systemic anti-LG antibody responses may serve as biomarkers for the identification of patients at increased risk for a distinct form of disease flares and progression of tissue injury. By elucidating the mechanisms through which intestinal RG expansions affect SLE disease activity, we may contribute to the development of tools that lead to more accurate prognoses and safer, targeted therapy.

In summary, our findings suggest that blooms of strains of a common gut commensal contribute to clinical flares in a subset of LN patients. This microbial gut expansion leads to impaired gut barrier function, exposing the host immune system to pathogenic microbial products and triggering subsequent systemic immune activation. The immune activation driven by gut dysbiosis in LN patients exhibits a similar, but lower-intensity profile of previously documented evidence of platelet activation and subsequent neutrophil degranulation, as that observed in patients acutely affected by bacterial sepsis. This study offers new insights into the mechanisms by which RG dysbiosis contributes to LN pathogenesis, highlighting the potential clinical utility of the anti-LG antibody as a surrogate marker for RG blooms. Together, these pathways contribute to the identification of a unique, previously unsuspected thromboinflammatory pathologic endotype in a subset of patients with proliferative LN.

## Methods

### Ethics Statement

The study was conducted according to the Declaration of Helsinki. Written informed consent, approved by the New York University Institutional Review Board (NYU IRB), was obtained from all subjects before their inclusion in the study for research use and publication of their data.

### Study Design (Human Subjects)

Patients were consecutively recruited from the NYU Langone Medical Center and Bellevue Hospital. All patients fulfilled the American College of Rheumatology (ACR) criteria for diagnosing systemic lupus erythematosus (SLE)^18^. Patients were excluded from the study based on the following criteria: pregnancy or breastfeeding, recent or current serious confounding medical disorders, current malignancy other than skin cancer, or cyclophosphamide treatment within the last 12 months. Doses of azathioprine, mycophenolate mofetil (MMF), or methotrexate were unchanged for the 4 weeks before study entry. Within the last 3 months, no patient had a history of serious infection requiring hospitalization, nor antibiotic treatment.

### Patient Cohorts

We evaluated a pilot cohort consisting of 8 healthy controls and sixteen SLE patients (eight with renal disease and eight without) from NYU. All lupus nephritis (LN) patients had class III and/or IV disease by renal biopsy, with elevated dsDNA, hypocomplementemia, but varied based on the presence of rash and arthritis. All patients received daily hydroxychloroquine, either alone or in combination with methotrexate or azathioprine, and one patient in each group received MMF. Additionally, 3 out of 4 patients in each group were on prednisone at doses ranging from 20 to 40 mg (**Extended data table 1)**. No patients with anti-phospholipid antibody (APL) positivity were included in the pilot cohort. Our validation cohort was subjects from the ACCESS_LN study who received only standard of care treatment^23^. From this validation cohort, data on fecal microbiota communities were unavailable. At baseline, all patients had a urine protein-to-creatinine ratio (Pr/Cr) greater than one and were experiencing a renal disease flare at the time of enrollment. Patients had not received prior treatment with cyclophosphamide or azathioprine. No patient received 40 mg or higher daily doses of prednisone. There was no significant difference in available APL antibody levels between patients with positive versus negative anti-LG antibody levels **(Extended data table 2)**. Publicly available data for meta-analysis of platelet and neutrophil. Interferon, and B cell whole-blood gene expression signatures included data from Wither et al. 2018^19^ (GSE99967, n =11, Canada, PAXgene blood collection, Microarray), Figgett et al. 2019^20^ (GSE112087, n=15, Australia, PAXgene blood collection, RNAsequencing), Chiche et al. 2014^21^ (GSE49454, n=64, France, Tempus tubes, Microarray), and Banchereau et al. 2016^22^ (GSE65391, n=331, USA, Tempus tubes, Microarray), and only samples with noted meta data available for active renal disease were included. Both male and female patients were included. Data were normalized through Z-score normalization across all SLE sample gene expression data in each experiment, and then all identified active LN samples were included in spearman correlation analysis completed.

### Mice

All murine procedures in this study adhered to protocols approved by the Institutional Animal Care and Use Committee (IACUC) at NYU Grossman School of Medicine. Female C57BL/6 and platelet/megakaryocyte reporter mice (TdTomato-Gp1bCre)^30^ were locally bred under specific-pathogen-free (SPF) conditions and were aged between 8 and 12 weeks at the commencement of experimentation. All mice used in the experiments were female. Additional studies were completed in FcγRIIb^-/-^ mice, a well-established lupus-prone model^28^ due to the absence of the sole inhibitory Fcγ-receptor on B cells, which leads to accelerated systemic autoimmunity.

### Intestinal colonization of *Ruminococcus gnavus*

Mice aged 8 to 12 weeks were preconditioned with an oral antibiotic cocktail comprising ampicillin (1g/L; Fisher Scientific) and enrofloxacin (0.5g/L, Sigma Aldrich) for two weeks, plus an additional four days of a triple-antibiotic cocktail including ampicillin (1g/L, Fisher Scientific), enrofloxacin (0.5g/L, Sigma Aldrich), and metronidazole (1g/L, Fisher Scientific). The antibiotic solutions were freshly prepared weekly in autoclaved drinking water using a magnetic stirrer for approximately twenty minutes to ensure complete solubility. Antibiotic treatment was administered in 300 mL clear glass bottles supplied by NYU Langone. After antibiotic pre-treatment, C57BL/6 and TdTomato-Gp1bCre mice were colonized with an LG-producing RG strain isolated from an SLE patient with an RG bloom, as previously reported^7,25^. FcγRIIb^-/-^ mice were colonized with RG strains with (nb92, RG2) or without (nb90) lipoglycan. Mice were orally gavaged with 8 × 10^7 CFU of bacteria diluted in sterile PBS or sham gavaged, for a total of five administrations, over 2 weeks.

Intestinal permeability assay was performed as described previously^11,25^. To evaluate intestinal permeability, mice underwent a 4h fasting period before the experiment. After fasting, they received an oral gavage of 4,000-Da fluorescein isothiocyanate (FITC)-dextran (FD4) (Sigma-Aldrich, St. Louis, MO) at a dosage of 250 mg/kg body weight in 200 μL of buffered saline.

Blood samples were taken via cheek bleed via a lancet after 3h. The concentration of FD4 in the serum was measured using a fluorimeter set at an excitation wavelength of 490 nm and an emission wavelength of 530 nm. Serum samples were serially diluted to create a standard curve for quantifying FD4 concentration. An increase in serum fluorescence intensity signified enhanced intestinal permeability.

### Intraperitoneal challenge with LG preparation

Lipoglycan, from RG strains isolated from patients with SLE during disease flares, was purified by hydrophobic interaction chromatography as described earlier^1,7^. A single intraperitoneal (IP) injection of 1.5-2 μg/g of purified LG was administered to five-to seven-month-old wildtype female mice. Six days post-injection, mice were sacrificed, and spleen tissues were collected for histological assessment.

### Quantitative PCR analysis

Fecal pellets were collected before and 21 days after cessation of antibiotic treatment. Fecal genomic DNA was extracted using the DNeasy PowerSoil Pro Kit (Qiagen) following the manufacturer’s protocol, and DNA quantified using a Nanodrop 1000 instrument (Thermo). Each qPCR reaction contained a 25 μL sample consisting of 9.5 μL molecular grade water, 0.25 μL of FWD-Primer at 100 μM, 0.25 μL of REV-Primer at 100 μM, 10 μL of SYBR Green PCR Master Mix (Applied Biosystems), and 5 μL of a sample containing 15 ng of total genomic DNA.

Total bacterial 16S rRNA quantitation was assessed using the following primers:

UniF340 (5’-ACTCCTACGGGAGGCAGCAGT-3’)

UniR514 (5’-ATTACCGCGGCTGCTGGC-3’)

1. *R. gnavus* species-specific 16S rRNA quantitation was assessed using the following primers:

FWD (5’-GGACTGCATTTGGAACTGTCAG-3’)

REV (5’-AACGTCAGTCATCGTCCAGAAAG-3’), as described previously^11,25^.

The qPCR assay was run on the StepOnePlus™ Real-Time PCR System (Thermo Fischer). Thermal cycling conditions were as follows: initial denaturation at 95°C for 5 minutes, followed by 40 cycles of 95°C for 15s and 58°C for 30s. The melting curve stage included 95°C for 15s, 60°C for 1min, and 95°C for 15s. qPCR confirmed a >100-fold decrease in total 16S rRNA in all mice, indicating sufficient depletion of the intestinal bacterial burden.

### Flow cytometry and Platelet activation stimulation assays

Bone marrow cells from the whole blood and femur were collected from platelet/megakaryocyte reporter mice (TdTomato^fl/fl/fl^ Gp1b-Cre+) or C57BL/6 mice. Diluted whole blood (1:20 in modified Tyrode’s s-HEPES buffer with 3% BSA) was used for platelet integrin activation assays, staining for CD41+ (Miltenyi Biotec) and P-selectin (BD Biosciences) in B6 mice and TdTomato^fl/fl/fl^ Gp1b-Cre+ reporter mice. The samples were then stimulated with native collagen fibrils (Type I from equine tendons suspended in isotonic glucose, 5µg/mL, Chrono-Log), adenosine diphosphate (ADP, 10 µM, Chrono-Log), or thrombin from human plasma (0.1 U/mL, Chrono-Log) for 15 min. Bone marrow cells were then stained with Zombie Yellow™ dye (1:100) in 1X PBS at a concentration of 1 × 10^6 cells per 100 μL for 20 min at room temperature in the dark. Following this, the cells were stained with CD41+ (Miltenyi Biotec) and CD61+ (Miltenyi Biotec) antibodies to detect bone marrow megakaryocytes. Samples were analyzed using an Attune NxT flow cytometer (Thermo Fisher Scientific) or an ID 7000 spectral flow cytometer (Sony), and the acquired data were processed using FlowJo software (FlowJo v10.10, LLC).

### Anti-Lipoglycan (Anti-LG) antibody measurement

Serum samples were collected in endotoxin-free vacutainer tubes (BD Biosciences) without anticoagulant. Anti-LG antibody levels were measured using a bead-based standardized assay with a monoclonal anti-LG internal control with a positive cut-off defined as >53,000 U/mL. Longitudinal measurements were taken up to 105 weeks, as described earlier^7^.

### Platelet Factor 4 and Neutrophil Extracellular Trap (NET) Fragment ELISA

Serum platelet factor 4 (PF4) levels were measured using the PF4 Quantikine ELISA Kit (R&D Systems). NET fragments were quantified using a multiplex ELISA that combines antibodies against myeloperoxidase (MPO), citrullinated histone H3 (CitH3), and DNA^31^, with units (AU) based on interpolation of 450nM absorbance to a serial dilution standard curve of supernatant from phorbol myristate acetate stimulated, purified neutrophils, as described^32^.

### Histologic studies and multiplex IHC

Spleen tissues were fixed for 48h in 10% neutral buffered formalin and stored long-term in 70% ethanol. Formalin-fixed, paraffin-embedded (FFPE) samples were serially sliced into 5 μm-thick cross-sections at the NYU Experimental Pathology core facility at NYU. Immunofluorescence (IF) staining for megakaryocytes was performed using antibodies against CD41, DAPI, and CD19. Large cells (>20uM diameter) with CD41+ staining on the cell surface were counted as megakaryocytes, while smaller CD41+ fragments were considered platelets and therefore not quantified. Quantification was completed based on the number of cells per high-powered field at 40x magnification for all sections, with 4 images per spleen taken for experiments, with at least three mice per group in all experiments shown. Hematoxylin and eosin (H&E) quantitative assessments were completed blindly (AA, SG). Quantification of total prominent lymphoid aggregates was scored in equivalent total length distal small intestine swiss rolls, with scoring weight (range 2+ to 10.5+) determined based on the number of lymphoid cells per aggregate.

Scoring system included +0.5 for aggregates with 25-55 lymphocytes, +1 if 56-149 count aggregate, +2 for 150–299 count aggregate, +3 for 300+ count per aggregate, and +1 additional point if a germinal center was distinctly identifiable in the lymphoid aggregate. For renal H&E morphologic evaluation of FcγRIIb^-/-^, determination of the activity index was completed with an activity index scoring system^33^, which included assessment for mononuclear cells in the tubulointerstitial, leukocyte infiltration (in glomeruli), endocapillary hypercellularity, and hyaline thrombi/basement membrane splitting/early crescent formation. Each component was scored 0-

For multiplex IF and imaging processing, FFPE sections were stained with Akoya Biosciences® Opal™ multiplex automation kit reagents (Leica, Cat. # ARD1001EA) on a Leica BondRX® autostainer, according to the manufacturer’s instructions. In brief, slides underwent sequential epitope retrieval with Leica Biosystems Epitope Retrieval, primary and secondary antibody incubation, and tyramide signal amplification with Opal® fluorophores. Primary and secondary antibodies/polymers were removed during sequential epitope retrieval steps while the relevant fluorophores remained covalently attached to the antigen, as shown in Extended Data Table 4. Slides were counterstained with spectral DAPI (Akoya Biosciences, FP1490) and mounted with ProLong Gold Antifade (ThermoFisher Scientific, P36935). Semi-automated image acquisition was performed on a Vectra® Polaris multispectral imaging system. For colonization IF experiments, samples from sham-vs. the nb92 substrain-, and the nb90 substrain- vs. RG2- colonized mice, were included on the same slide, stained, and quantified in parallel.

### Whole Blood RNA Sequencing

For human samples, whole blood was directly collected into RNA stabilizing PAXgene tubes and extracted with the standard RNA extraction protocol (Qiagen). Unprocessed FASTQ RNAseq data for validation ACCESS_LN cohort^23^ provided by the Immune Tolerance Network^34,35^. Whole blood samples from mice were obtained through retro-orbital bleeding and mixed with 0.1M sodium citrate (blood ratio = 10:1). RNA was extracted using a modified protocol of the PAXgene Blood RNA Kit (Qiagen, Valencia, CA, USA), originally designed for human whole blood^36^. All RNAseq data were pre-processed with the same methodological pipeline, in parallel, through the NYU Langone Genome Technology Center Core.

### Data Preprocessing

FASTQ files from RNA sequencing were processed using the Seq-N-Slide pipeline. The quality of reads was assessed using FASTQC, FastQ Screen, and Picard tools. Samples were then aligned to the human genome (hg38) using the STAR aligner, and reads were quantified using featureCounts. All samples used in this study consisted of high-quality RNA-seq data, each exceeding 1,000,000 read counts. For longitudinal pilot cohort data, samples were run in two separate runs, including exact technical replicates in both experiments to ensure control for batch effects. A stringent gene expression filter was applied, in which genes exhibiting consistently low expression levels (less than four counts) were removed, retaining only those transcripts with a count of at least four across all samples. This pre-filtering approach corrected for any batch effects between runs (Extended data 13).

### Bioinformatic, Statistical Analyses, and Visualization

All bioinformatic and statistical analyses were performed using R. Differential gene expression analyses were conducted using DESeq2. Gene set enrichment analysis (GSEA) was performed to identify significantly enriched pathways. This method compares the expression levels of predefined gene sets between different conditions to determine if specific pathways are overrepresented in the data, providing insights into the underlying biological processes. Genes within pathways were defined by Gene Ontology (GO) or Reactome databases. Modules were defined through the SingScore package in R, which ranks gene expression values and calculates a normalized score that reflects the relative expression of the gene set within each sample. The genes included for the platelet module set are MMRN1, SNCA, SPARC, CD109, PF4, PROS1, EGF, PDGFA, F13A1, ABCC4, GATA1, MPP1, MYL9, MEIS1, VWF, P2RY12, RAB27B, PF4V1, and ITGB3. For the neutrophil genes module: SLC7A11, VTI1B, STOM, LGALS3, PGRMC1, SLC18A2, CLU, ADAM9, THBS1, CD300LF, MILR1, SNAP23, IFNGR2, STXBP3, PIK3CD, and SBNO2. The B cell module genes included CD27, GCSAM, NOD2, IGHG1, and MZB1, and the IFN module genes included IFI44, IFI44L, and RSAD2. Statistical analyses were completed within the R environment, or with Graphpad Prism V10. Figure infographics produced with Biorender.

## Acknowledgements

This study was in part funded by NIH P50-AR070591, R01 AI143313, R21AI180737, and the Lupus Research Alliance (GJS), as well as through the Lupus Foundation of America Gilkeson Career Development Award. (AA). AA is supported in part by NIH 5T32AR069515-07. We are grateful to the Immune Tolerance Network, Laura Cooney, and Sharon Chung. Histological analyses were completed through the NYU Grossman School of Medicine Experimental Pathology Core, including Mark Alu and Gyles Ward. RNA sequencing studies were completed through the NYU Genome Technology Center, supervised by Dr. Adriana Heguy. Flow cytometry studies were completed through the Cytometry and Cell Sorting Laboratory, with assistance from Catherine Rapelje, Florenal Joseph, and Jiangwei Li. These NYU facilities are partially funded by the Cancer Center Support Grant NIH/NCI 5 P30CA16087 at the Laura and Isaac Perlmutter Cancer Center. We thank Peter Izmirly and Jill Buyon for their clinical support. We are grateful to Kate Trujillo, Uzair Chaudhary, Steven Medvedovsky, and Heiko Käßner for their technical assistance; Bharati Matta and Tatiana Borja for their advice on neutrophil assays; and assistance with platelet activation flow cytometry assays. We are grateful to Marc Scherlinger, Patrick Blanco, and Eric Morand for their discussions. We thank all our patients for their participation, and the clinical coordinators in this study.

## Author Contributions

Study design: GJS and AA. Flow cytometric analysis: AA, CFR. RNA sequencing analyses: AA, MC, KR. Platelet and neutrophil immunoassay methodology, completion, and analyses: AA, AL, TY, BB. Histology analysis: AA, SG, CL. Murine model management and colonization studies: AL, ZA, MY, CFR, AA, GJS. *R.gnavus* characterization, maintenance, and bacterial pellet generation: NU, GJS. Lipoglycan isolation: NG. Platelet activation studies: AA, CFR, MC, JP, BR, KR. Figure formulation: AA and GJS. Manuscript writing AA, GJS. Manuscript editing GJS, AA, CFR, NG, AL, TW, ZA, BR, KR.

## Data availability statement

Clinical data for patients in our pilot cohort are available in our prior publications^1,7^, and for the validation cohort included in extended data, plus through the Immune Tolerance Network^23^ (itntrialshare.org) of NIAID. Datasets for RNA sequence data will be made available in online repositories. Additional primary data are available on request from qualified parties.

