## Supplemental Figures, Tables, and Legends for "Pathogenic strains of a gut commensal drive systemic platelet activation and thromboinflammation in lupus nephritis"

**Extended Data Table 1.** Clinical characteristic summary of eight lupus nephritis patients in the initial test pilot whole-blood RNAseq cohort.

| Patient ID | Age | Ethnicity | SLEDAI | Medications | RG Bloom vs. without bloom |
|---|---|---|---|---|---|
| s47 | 39 | Asian | 8 | MMF<br>AZA<br>Prednisone 5mg | RG Bloom |
| s78 | 38 | White Hispanic | 8 | MMF<br>Prednisone 5mg | RG Bloom |
| s107 | 32 | White Hispanic | 22 | Prednisone 20mg | RG Bloom |
| s120 | 37 | Black | 15 | Prednisone 40mg | RG Bloom |
| s89 | 37 | Asian | 6 | Prednisone 40mg | Without Bloom |
| s124 | 35 | White Hispanic | 16 | MMF<br>Prednisone 20mg | Without Bloom |
| s172 | 24 | Asian | 12 | MMF | Without Bloom |
| s202 | 38 | White | 2 | MMF | Without Bloom |

**Extended Data Table 1.** AZA, azathioprine; MMF, mycophenolate mofetil; Oral steroid dose noted represents daily prednisone equivalents.

**Extended Data Table 2.** Summary of Pathway Analysis Genes and Overlapping Genes. This .xlsx table highlights the genes identified through pathway analysis of top 20 pathways enriched in LN RG patients vs. LN patients without dysbiosis in the pilot cohort. Of these top 20, 5 related to platelet activation or degranulation.

**Extended Data Table 3.** Demographics of validation cohort.

| **Demographics** | **Anti-LG Negative** | **Anti-LG Positive** | **Sig** (p) |
|---|---|---|---|
| **Count** | N (%) | N (%) | |
| **Sex** | 8 | 12 | 0.40 |
| Male | 0 (0) | 1 (8.3) | |
| Female | 8 (100) | 11 (91.7) | |
| **Race/Ethnicity** | 8 | 12 | 0.62 |
| Black | 3 (37.5) | 6 (50) | |
| Hispanic | 1 (12.5) | 0 (0) | |
| Asian | 1 (12.5) | 1 (8.3) | |
| White | 3 (37.5) | 5 (41.7) | |
| **Age** | 8 | 12 | 0.25 |
| 18 – 24 years | 4 (50) | 2 (16.7) | |
| 25 - 39 years | 3 (37.5) | 6 (50) | |
| >= 40 years | 1 (12.5) | 4 (33.3) | |
| **Age** (years, median, 95% CI) | 24.5 (19, 48) | 35.0 (28, 46) | **0.02** |
| **Lymphocyte Count** (x10^3/uL) | 1.56 (0.7,4.9) | 1.48(0.66,2.0) | 0.57 |
| **Platelet Count** (x10^3/uL) | 248 (159,334) | 290.5 (224,305) | 0.73 |
| **Urine Pr/Cr Ratio** (g/g) | 2.65 (6.1,7.5) | 1.6 (1.2, 5.3) | 0.18 |
| **eGFR** | 78.6 (18.3,109.6) | 65.6 (41.1, 85.0) | 0.57 |
| **Albumin** | 2.65 (1.2, 4) | 2.9 (2.3, 3.4) | 0.35 |
| **BILAG** (Non-Renal Total Score) | 3 (1, 15) | 3 (1, 6) | 0.61 |
| **Immunomodulatory Medication Use** | 8 | 12 | |
| Hydroxychloroquine | 7 (87.5) | 8 (66.7) | 0.29 |
| Azathioprine | 2 (25) | 1 (8.3) | 0.31 |
| Mycophenolate Mofetil | 4 (50) | 6 (50) | 0.99 |
| **Pred Equiv at Baseline** (mg) | 8 | 12 | 0.33 |
| None | 2 (25) | 3 (25) | |
| 10-20 mg | 1 (12.5) | 5 (41.7) | |
| 21-30 mg | 3 (37.5) | 1 (8.3) | |
| >30 mg | 2 (25) | 3 (25) | |
| **Lupus ISN/RPS Classification** | 8 | 12 | 0.40 |
| III Only | 1 (12.5) | 4 (33.3) | |
| IV Only | 1 (12.5) | 2 (16.7) | |
| III + V | 3 (37.5) | 1 (8.3) | |
| IV + V | 3 (37.5) | 5 (41.7) | |
| **Anti-Coagulant Use** | 8 | 12 | |
| Aspirin | 1 (12.5) | 5 (41.7) | 0.20 |
| Warfarin | 0 (0) | 1 (8.3) | 0.42 |
| Heparin | 1 (12.5) | 0 (0) | 0.20 |
| **Anti-Cardiolipin antibody** | 8 | 12 | 0.29 |
| Positive | 1 (12.5) | 4 (33.3) | |
| **dsDNA** (IU/mL) | 8.5 (0, 911) | (0, 200) | 0.50 |
| **C3** (mg/dl) | 79 (28, 94) | (48, 102) | 0.75 |
| **C4** (mg/dl) | 13 (8, 22) | (10, 23) | 0.64 |

**Extended Data Table 3.** Demographics (sex, race/ethnicity, age), immunomodulatory medication use, anticoagulant use, anti-phospholipid antibody (anti-cardiolipin IgM or IgG), and prednisone equivalents at baseline. Admission immunosuppression notes anti-malarial use, plus additional medications used. P-values represent chi-square statistics for all comparisons, except for lymphocyte count, platelet count, dsDNA, C3, C4, Ur pr/cr ratio which were assessed as an independent, non-parametric mann-whitney U test shown as median and 95% CI. Age is provided as both a categorical and ordinal variable. mg = milligrams; equiv = equivalents; Pr/Cr = protein/creatinine ratio; BILAG =British Isles Lupus Assessment Group. All patients included were ANA+. No patients were on LMWH/Nadroparin, patients with only Class V LN disease, or those with Pred >40mg were not included. Activity index, chronicity index, anti-beta2glycoprotein and lupus anticoagulant data were not available.

**Extended Data Table 4.** Antibodies used for multiple IHC.

| Renal Antibody Panel | | | | | | | | | | | |
|---|---|---|---|---|---|---|---|---|---|---|---|
| Target | Vendor, Cat.# | Clone | Retrieval | RRID | Dilution | Secondary HRP Polymer | Vendor/ Cat # | Incubation Time [min] | Opal TSA Fluorophore | Vendor/Cat# | Fluorophore Dilution |
| Cit-H3 | Abcam, ab232939 | EPR20358-13 | ER1-20 | NA | 100 | Rabbit-on - Rodent HRP polymer | Biocare/ RMR622L | 10 | 570 | Akoya/FP1488001KT | 150 |
| MPO | R&D, AF-3667 | Poly | ER1-20 | AB_2250866 | 300 | Goat HRP polymer 2 step | Biocare/ GHP516 | 10+10 | 520 | Akoya/FP1487001KT | 200 |
| CD3 | CST, 78588S | E4T1B | ER2-20 | AB_2889902 | 800 | Rabbit-on - Rodent HRP polymer | Biocare/ RMR622L | 10 | 690 | Akoya/FP1497001KT | 150 |
| F480 | CST, 70076S | D2S9R | ER2-20 | AB_2799771 | 1500 | Rabbit-on - Rodent HRP polymer | Biocare/ RMR622L | 10 | 480 | Akoya/FP1500001KT | 200 |
| Spleen Antibody Panel | | | | | | | | | | | |
| Target | Vendor, Cat.# | Clone | Retrieval | RRID | Dilution Factor | Secondary HRP Polymer | Vendor/ Cat # | Incubation Time [min] | Opal TSA Fluorophore | Vendor/Cat# | Fluorophore Dilution |
| CD41 | Abcam, ab134131 | EPR4330 | ER2-20 | AB_2732852 | 2000 | Rabbit-on - Rodent HRP polymer | Biocare/ RMR622L | 10 | 620 | Akoya/FP1495001KT | 150 |
| CD19 | CST, 90176S | D4V4B | ER2-20 | AB_2800152 | 1000 | Rabbit-on - Rodent HRP polymer | Biocare/ RMR622L | 10 | 520 | Akoya/FP1487001KT | 150 |

**Extended Data Table 4.** Panel of antibodies used for CD19, MPO, CitH3, F4/80, and CD3 staining in kidney and CD41, CD19 antibody staining in spleen with Vectra Polaris Fluorescence Scanning.

**Extended data 1.**

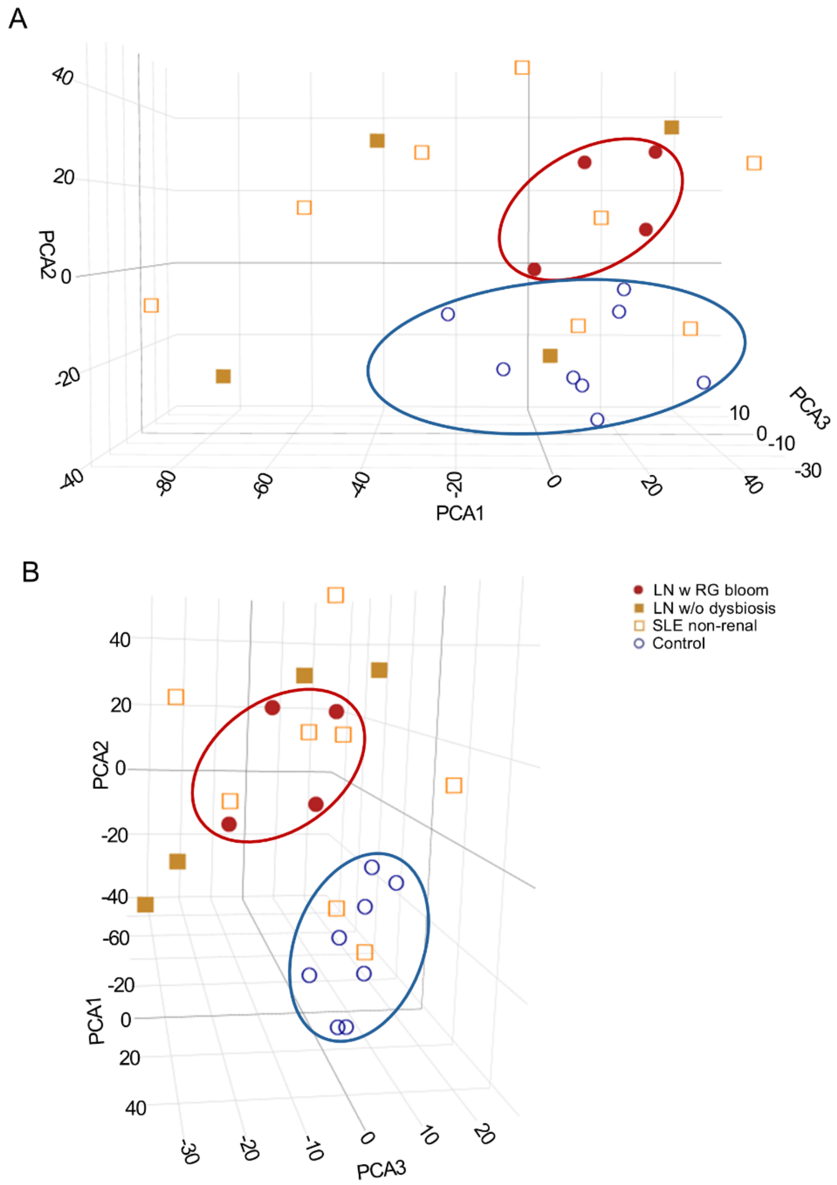


**Extended data 2.** Principal component analysis (PCA) of gene expression in whole-blood RNAseq libraries with top 25% variance among control, SLE without LN (SLE non-renal), LN without *R. gnavus* bloom (LN without dysbiosis), and LN with *R. gnavus* gut bloom (LN with RG bloom). Shown as two different visualizations of the same gene expression data, with PCA1, PCA2, and PCA3, representing 40.9%,19.1%, and 7.8% of the data, respectively.

**Extended Data 2.**

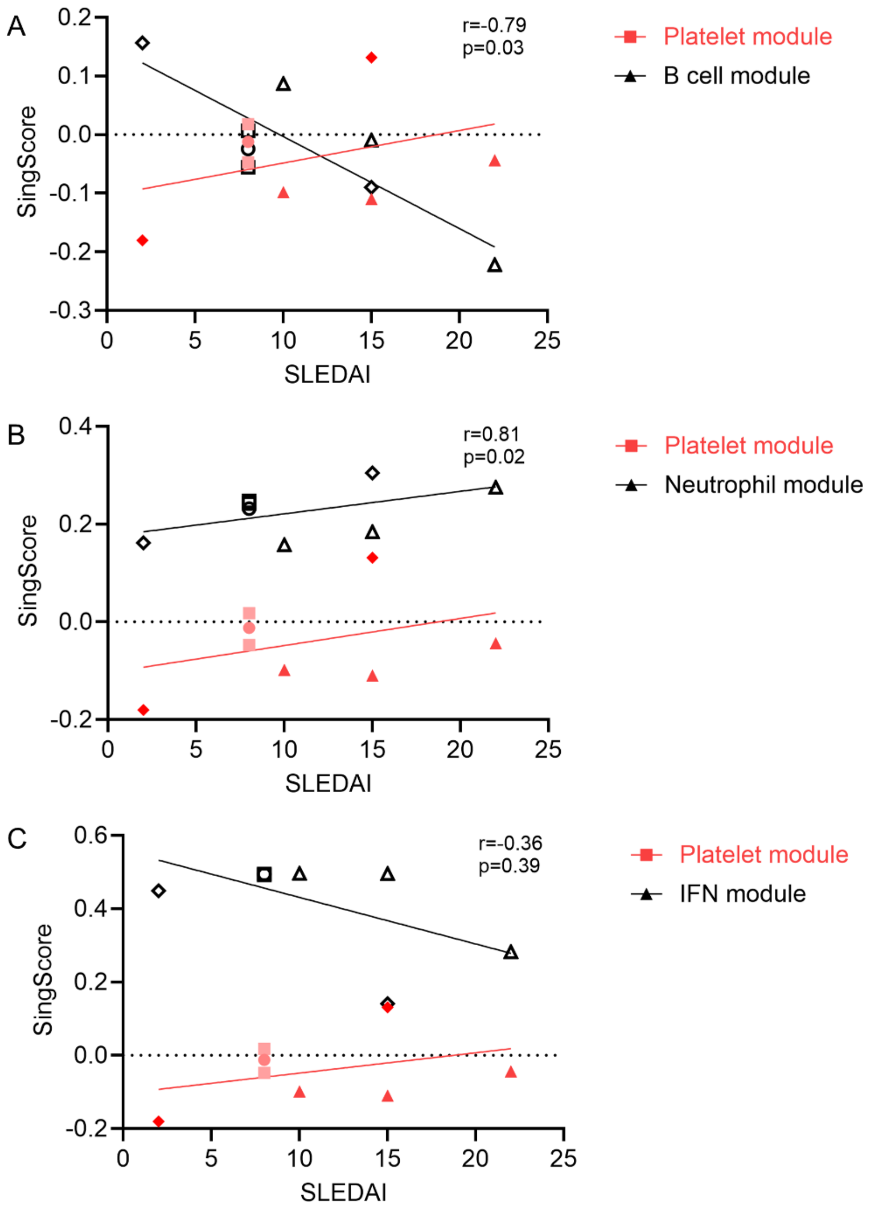


**Extended Data 2**. Longitudinal assessment of platelet vs. B cell activation genes. Gene-set modules (Singscore) of platelet activation vs. (A) B cell activity module, (B) neutrophil activation module, and (C) IFN gene modules in longitudinal whole blood RNAseq of patients with LN and gut dysbiosis (RG blooms). SLEDAI is noted as an indication of disease activity. Unique symbols represent the same patient in longitudinal measurements. Statistics represent spearman correlations.

**Extended Data 3.**

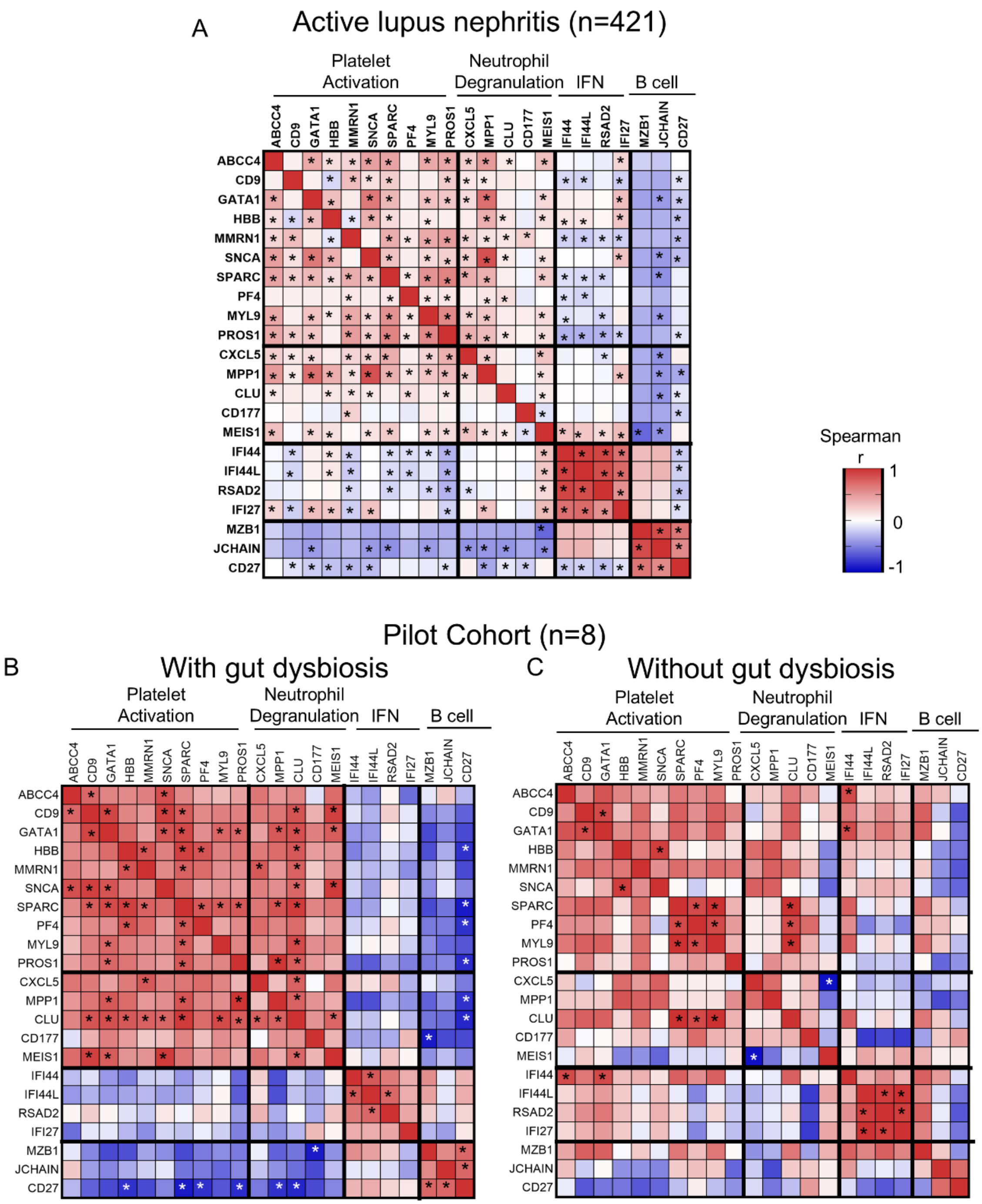


**Extended Data 3.** Correlation of platelet-related and IFN-related genes in LN patients. Spearman correlations of individual platelet, neutrophil degranulation, IFN stimulation, and B cell genes in whole-blood gene expression data (A) of 421 samples from patients with active lupus nephritis (from GSE99967, GSE112087, GSE49454, and GSE65391), (B) test cohort patients with gut dysbiosis and RG blooms (n=4 patients, 8 samples), and (C) test cohort patients without gut dysbiosis (n= 4 patients, 6 samples). * indicates $p<0.05$ for spearman correlation, showing overall an inverse correlation of platelet activation/neutrophil genes and IFN/B cell genes in patients with active LN overall, and more so in patients with gut dysbiosis than those without.

**Extended Data 4.**

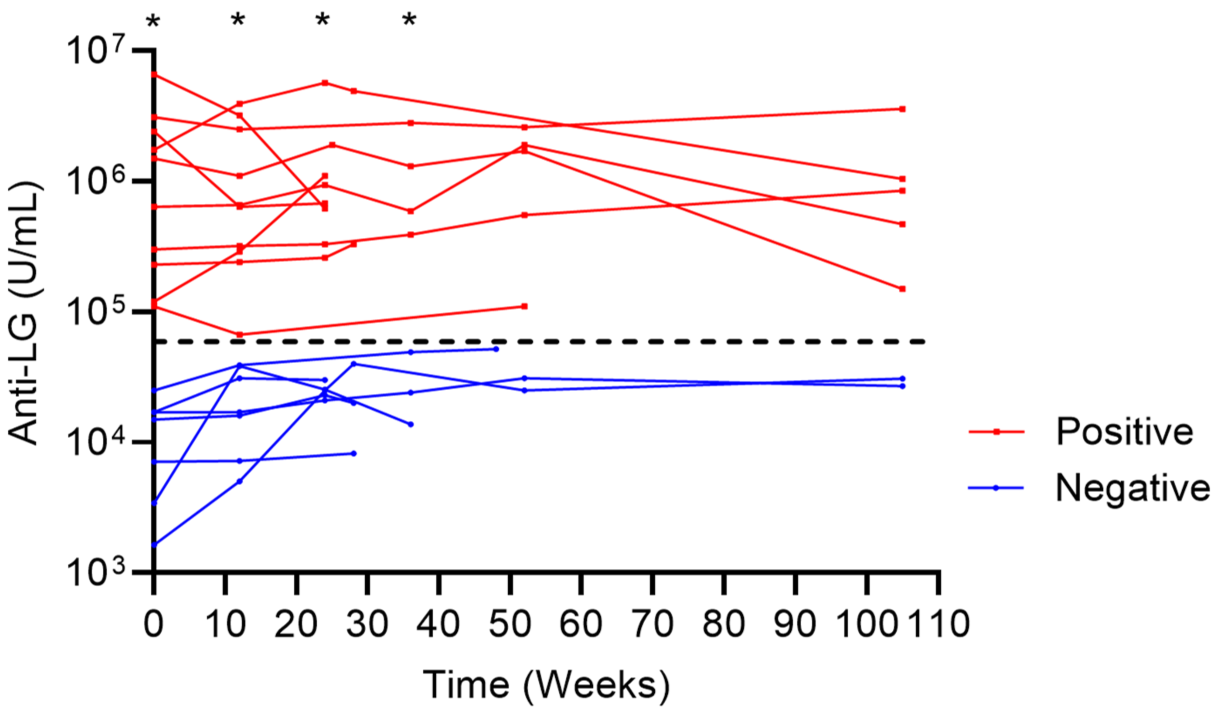


**Extended Data 4.** Stability of IgG against RG lipoglycan in LN patients overtime. Anti-lipoglycan antibody levels in patients designated as atni-LG positive or negative (based on a cut-off of 5.3E4 U/mL) measured overtime. *indicates $p<0.05$ by non-parametric, Mann-whitney U at study visit weeks 0, 12, 28, and 36 (+/- 4 weeks for visit window).

**Extended Data 5.**

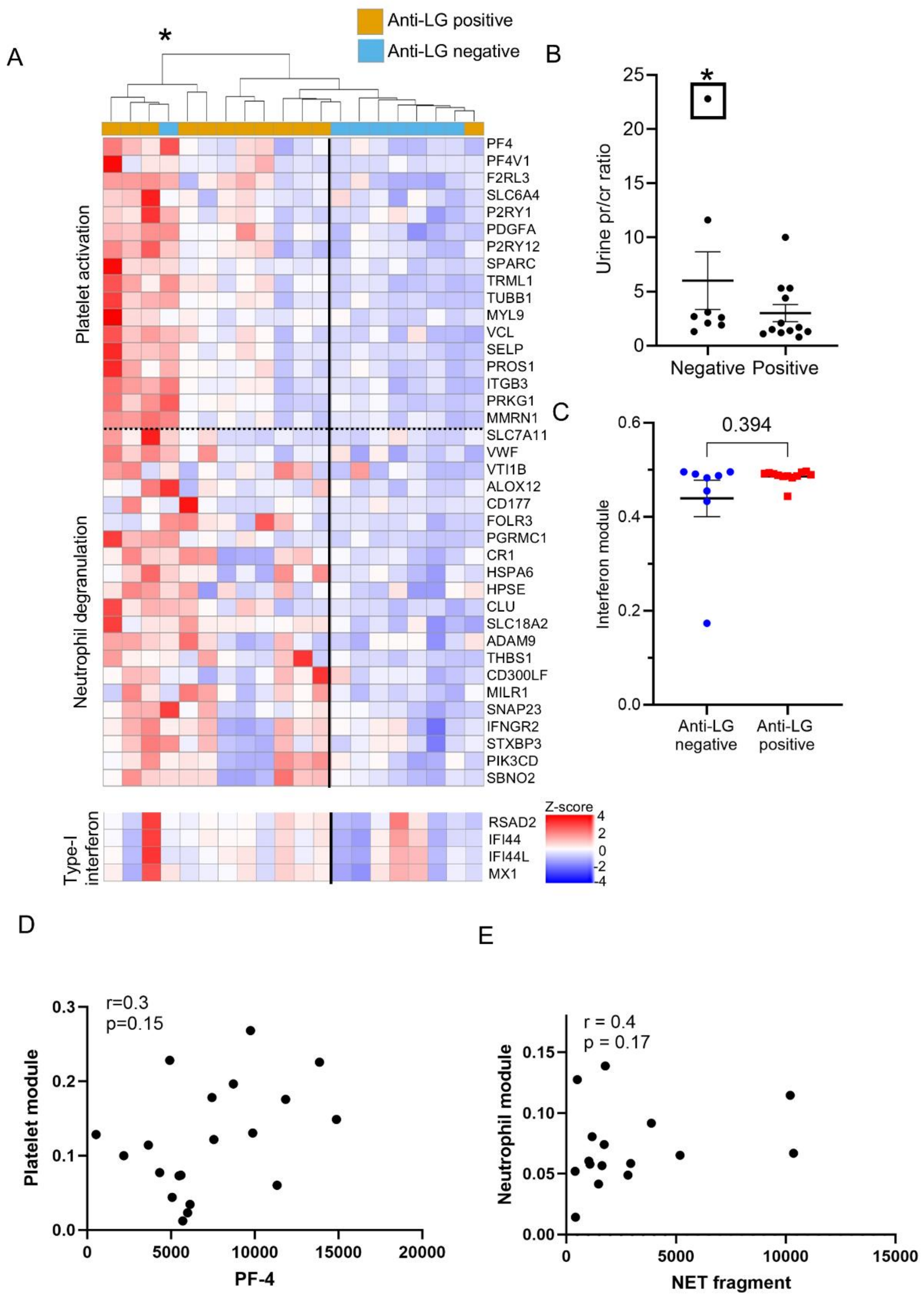

**Extended Data 5.** Unsupervised hierarchical clustering of validation cohort patients. (A) Clustering of platelet activation and neutrophil degranulation genes in validation cohort LN patients. Anti-LG positive patients (orange) cluster together, aside from one exception, and * indicates the one-anti-LG negative patient with high platelet activation gene signatures, who also has a Urine Pr/cr ratio >20 (B). (C) Type-I interferon genes are increased in both anti-LG positive and negative patients, with no significant difference in the IFN gene module. Unsupervised hierarchical clustering completed by Canberra linkage and median clustering. Statistics shown represent non-parametric t-test. (D,E) Spearman correlation r and p-value for direct correlation of (D) platelet gene module against PF-4 and (E) Neutrophil gene module against NET fragment (DNA, MPO, citH3) detection for patient samples in which whole-blood RNAseq and serum samples for ELISA testing were available.

**Extended data 6.**

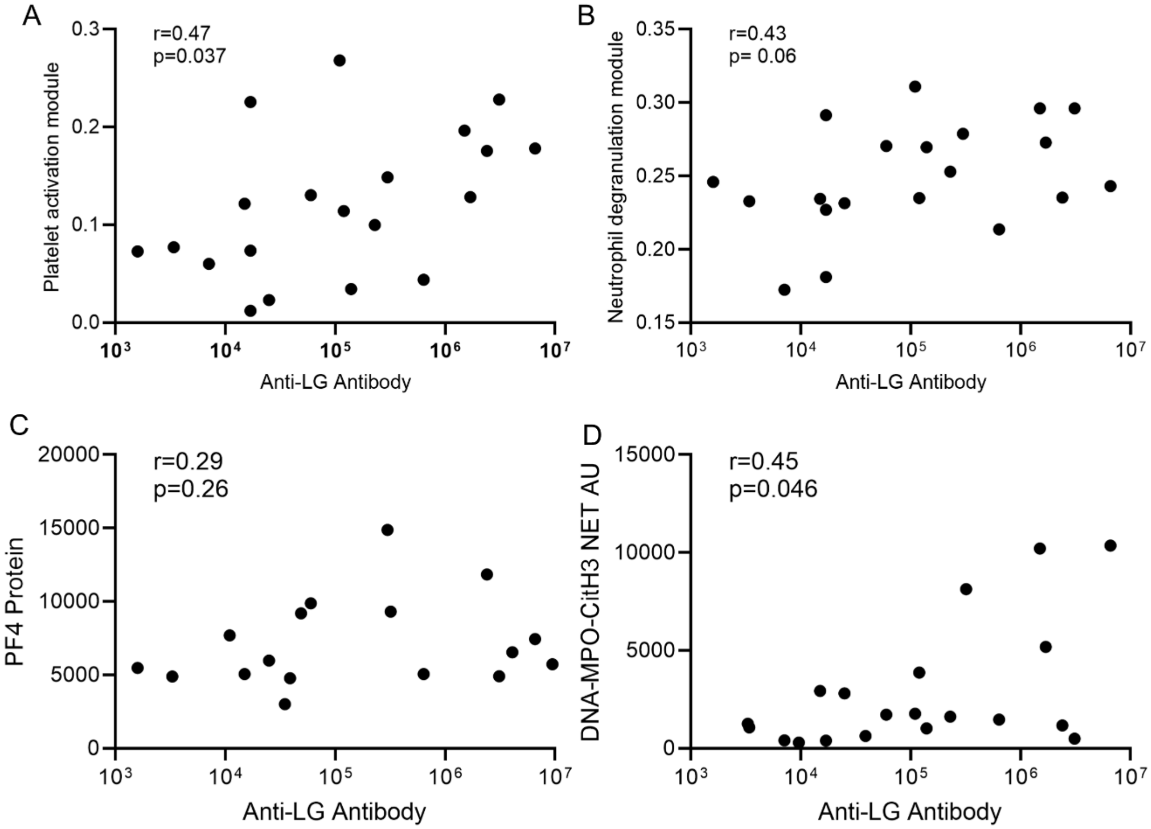


**Extended data 6.** Correlation of anti-LG antibody (U/mL) with platelet activation gene module (A), neutrophil degranulation gene module (B), platelet-factor 4 protein serum levels (ng/mL), and immunoassay-based detection of serum DNA-MPO-CitH3 complexes (NET fragments), representing evidence of NETosis activity (D). Statistics represent spearman r correlations.

**Extended Data 7.**

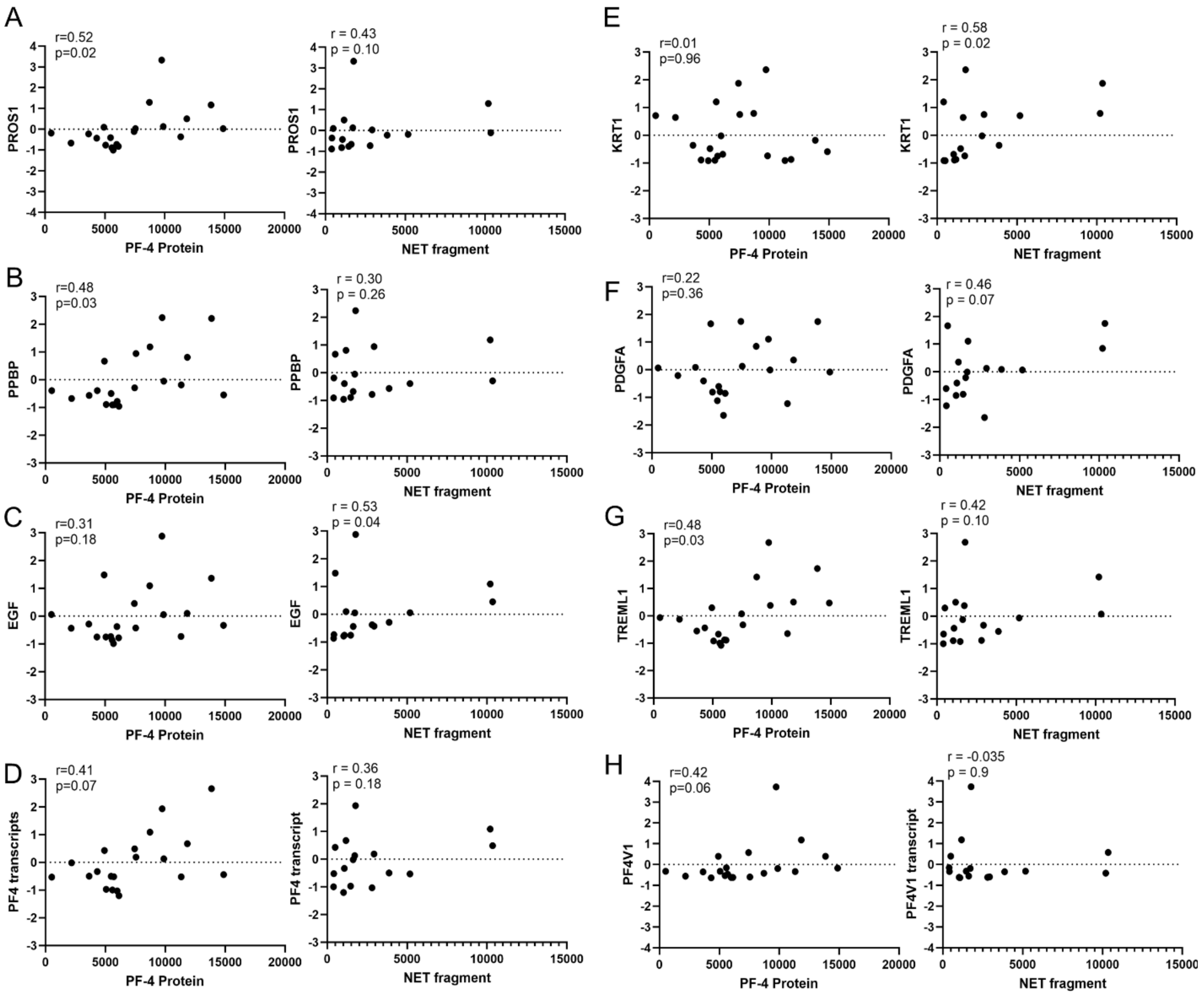


**Extended data 7.** Correlation of whole-blood RNAseq normalized transcripts against serum protein detection. Spearman correlation of Z-transformed normalized transcript counts of (A) PROS1, (B) PPBP (C) EGF, (D) PF-4, (E) KRT1, (F) PDGF-A, (G) TREML1, and (H) PF4-V1, compared to ELISA based detection in serum of PF-4 (Left) and DNA, MPO, citH3 positive NET fragments (Right).

**Extended Data 8.**

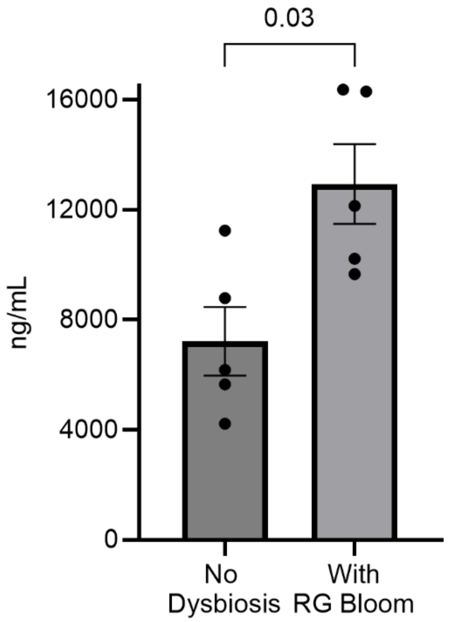


**Extended Data 8**. LPS binding protein in patients with and without RG blooms. Lipopolysaccharide (LPS) binding protein in patients with RG blooms vs. those without gut dysbiosis from our pilot cohort. P represents mann-whitney U, non-parametric test.

**Extended Data 9.**

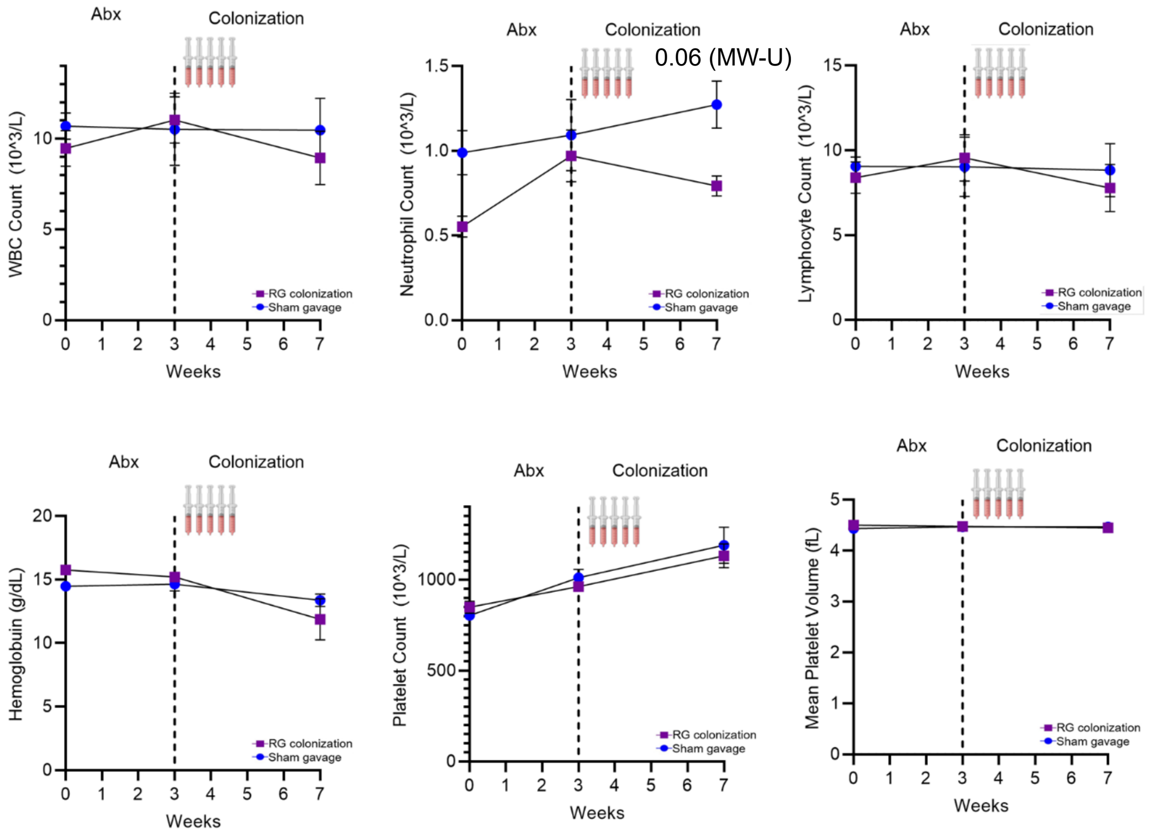


**Extended Data 9.** Colonization Sham vs. pathogenic lipoglycan producing RG strain (nb91). Complete blood count (CBC)-like analysis in B6 murine models of white-blood cells, neutrophils, lymphocytes, hemoglobin, platelet count, and mean platelet volume before the start of the experiment (week 0), after antibiotics (week 3, dashed line), and after peak RG colonization (week 7). Error bars represent mean with SEM, n = 7.

**Extended Data 10.**

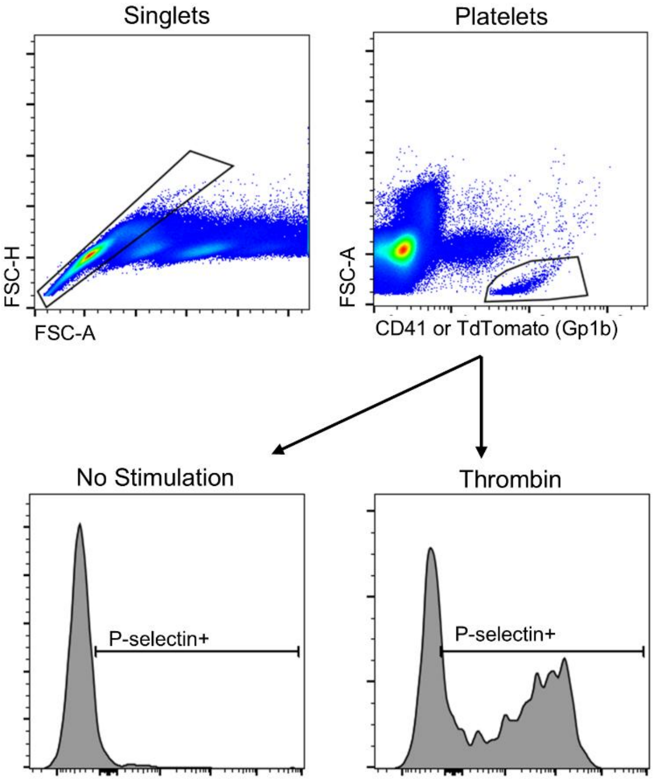


**Extended Data 10.** Gating strategy for activated platelets in whole-blood. Gating of singlets, platelets (Identified by CD41, or Gp1B), and subsequently P-selectin+ to define activated platelets. The same gating strategy was used for both C57/BL6 and TdTomato-Gp1bCre mice, comparing platelet activation with and without thrombin stimulation.

**Extended Data 11.**

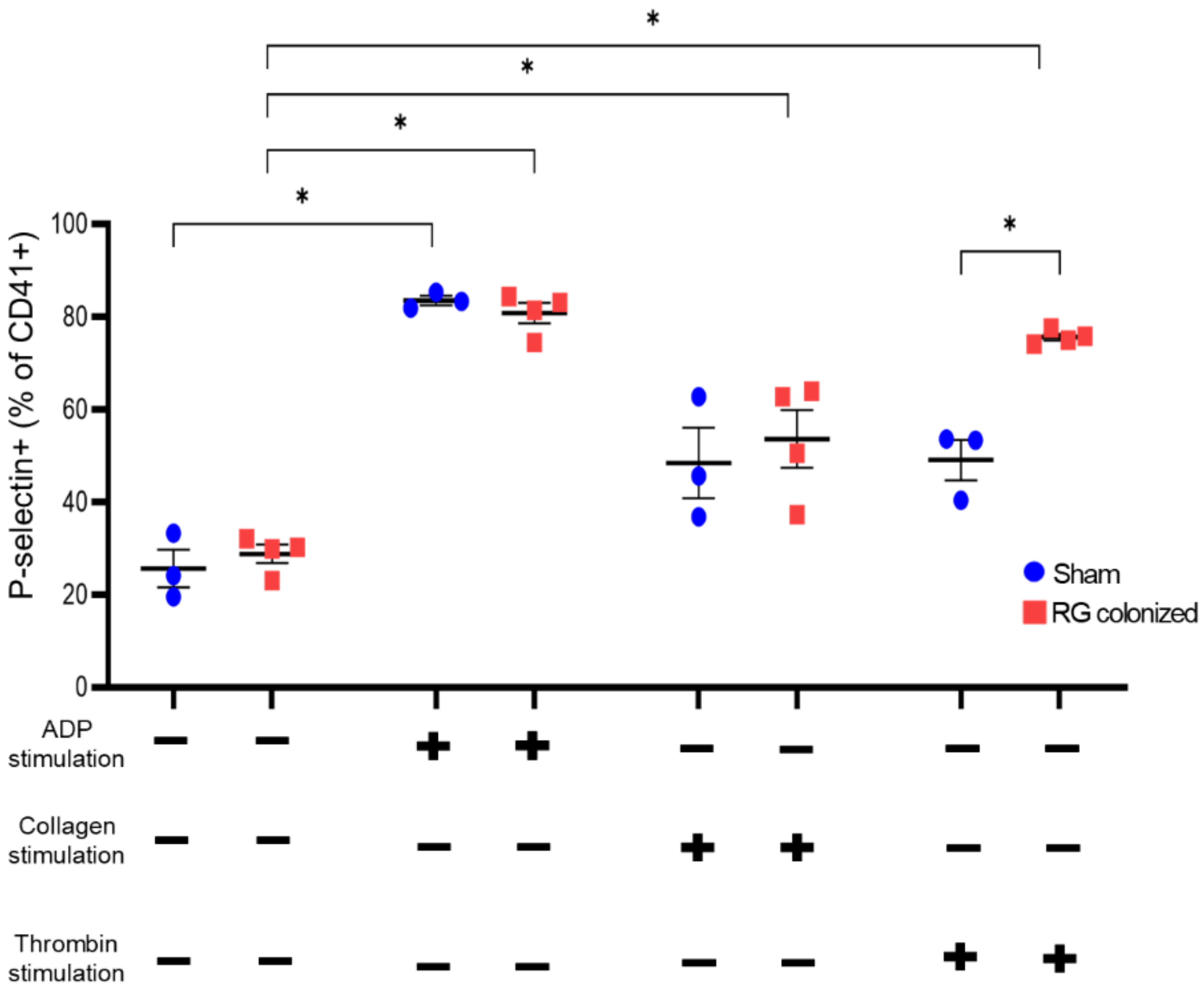


**Extended Data 11.** Platelet activation with Collagen and ADP. Platelet activation as measured by P-selectin surface expression after stimulation with lyophilized adenosine diphosphate or collagen fibrils (type I) from equine tendons in sham gavage and RG colonized mice. Unpaired t test, *p<0.05.

**Extended Data 12.**

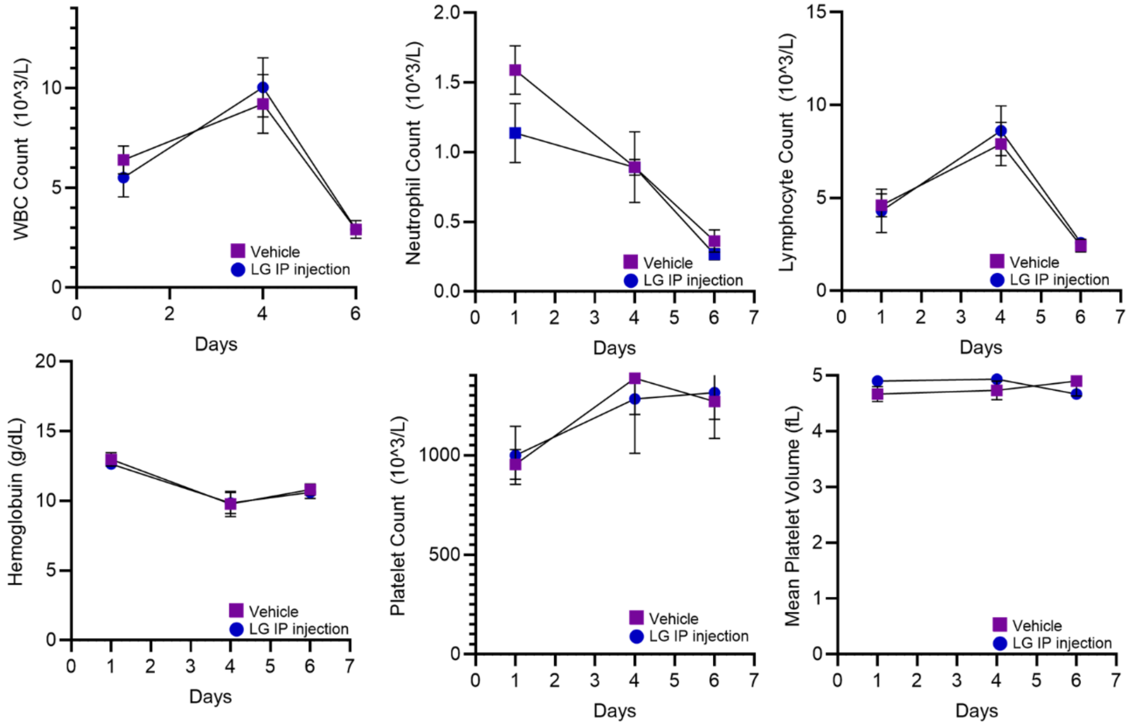


**Extended data 12.** Cell counts with Vehicle vs. lipoglycan injection. Complete blood count (CBC)-like analysis in FcγRIIb$^{-/-}$ lupus prone murine models of white-blood cells, neutrophils, lymphocytes, hemoglobin, platelet count, and mean platelet volume after 1,4 and 6, days after intraperitoneal (IP) injection of vehicle or lipoglycan (LG) in lupus-prone FcγRIIb$^{-/-}$ mice. 2ug/g of lipoglycan purified from nb91 was administered intraperitoneally at timepoint 0, and CBC data were collected at 8 hours, 4 days, and 6 days. Error bars represent mean with SEM. n = 6. No differences were also observed at 2 days post-LG intraperitoneal injection in another FcγRIIb$^{-/-}$ murine cohort.

**Extended data 13.**

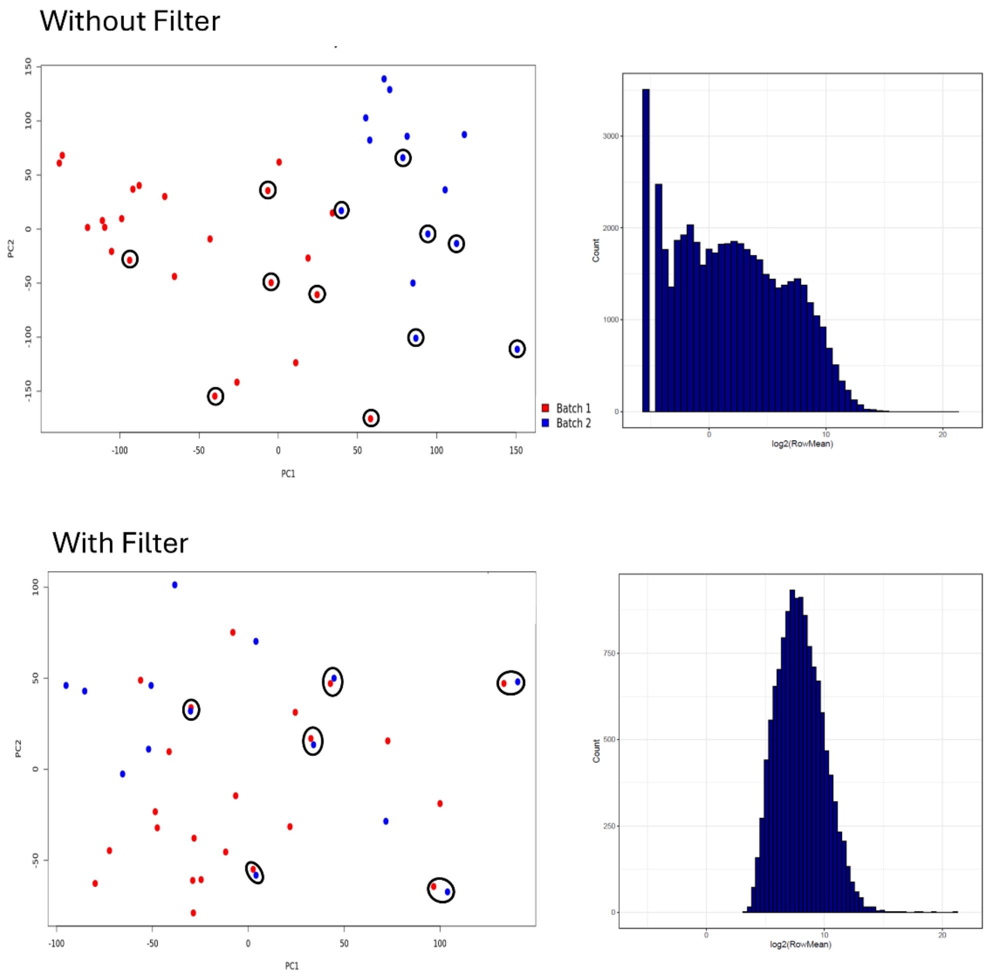


**Extended Data 13.** Filtering low-quality reads and batch effects assessment for pilot cohort RNAseq data. All samples used in this study consisted of high-quality RNA-seq data, each exceeding 1,000,000 read counts. To mitigate potential batch effects arising from the two RNA-seq runs, a stringent gene expression filter was applied. Genes exhibiting consistently low expression levels (less than 4 counts) were removed, retaining only those transcripts with a count of at least 4 across all samples. This filtering step reduced the initial 49,398 transcripts to 12,885 transcripts for subsequent analysis. Six technical replicates, representing all four experimental conditions (control, SLE without renal disease, LN without gut dysbiosis, and LN with RG bloom), were included in both RNA-seq runs to assess the impact of this filtering strategy. Principal Component Analysis (PCA) revealed clear batch effects in the unfiltered data. Following the removal of low-count genes, these batch effects were effectively eliminated, and technical replicates clustered tightly together. This demonstrates that our filtering strategy successfully removed noise and batch-specific variation, leading to more robust and accurate differential expression analysis using DESeq2, which incorporated both batch and condition (type) as factors in the design formula.
